# Variation in spatial population structure in the *Anopheles gambiae* species complex

**DOI:** 10.1101/2024.05.26.595955

**Authors:** Robert S. McCann, Jean-Paul Courneya, Martin J. Donnelly, Miriam K. Laufer, Themba Mzilahowa, Kathleen Stewart, Fiacre Agossa, Francis Wat’senga Tezzo, Alistair Miles, Shannon Takala-Harrison, Timothy D. O’Connor, Ag1000G Consortium

## Abstract

*Anopheles gambiae*, *Anopheles coluzzii*, and *Anopheles arabiensis* are three of the most widespread vectors of malaria parasites, with geographical ranges stretching across wide swaths of Africa. Understanding the population structure of these closely related species, including the extent to which populations are connected by gene flow, is essential for understanding how vector control implemented in one location might indirectly affect vector populations in other locations. Here, we assessed the population structure of each species based on a combined data set of publicly available and newly processed whole-genome sequences. The data set included single nucleotide polymorphisms from whole genomes of 2,410 individual mosquitoes sampled from 128 locations across 19 African countries. We found that *A. gambiae* sampled from several countries in West and Central Africa showed low genetic differentiation from each other according to principal components analysis (PCA) and ADMIXTURE modeling. Using Estimated Effective Migration Surfaces (EEMS), we showed that this low genetic differentiation indicates high effective migration rates for *A. gambiae* across this region. Outside of this region, we found eight groups of sampling locations from Central, East, and Southern Africa for which *A. gambiae* showed higher genetic differentiation, and lower effective migration rates, between each other and the West/Central Africa group. These results indicate that the barriers to and corridors for migration between populations of *A. gambiae* differ across the geographical range of this malaria vector species. Using the same methods, we found higher genetic differentiation and lower migration rates between populations of *A. coluzzii* in West and Central Africa than for *A. gambiae* in the same region. In contrast, we found lower genetic differentiation and higher migration rates between populations of *A. arabiensis* in Tanzania, compared to *A. gambiae* in the same region. These differences between *A. gambiae*, *A. coluzzii*, and *A. arabiensis* indicate that migration barriers and corridors may vary, even between very closely related species. Overall, our results demonstrate that migration rates vary both within and between species of *Anopheles* mosquitoes, presumably based on species-specific responses to the ecological or environmental conditions that may impede or facilitate migration, and the geographical patterns of these conditions across the landscape. Together with previous findings, this study provides robust evidence that migration rates between populations of malaria vectors depend on the ecological context, which should be considered when planning surveillance of vector populations, monitoring for insecticide resistance, and evaluating interventions.

## Background

Vector control interventions play a crucial role in preventing the transmission of malaria parasites and reducing the public health burden of malaria. These interventions have been associated with significant reductions in malaria parasite infection, uncomplicated and severe malaria burden, and all-cause child mortality [1–3]. In 2022, national malaria programs distributed 254 million long-lasting insecticidal nets (LLIN) and protected 62 million individuals through indoor residual spraying (IRS) according to the World Health Organization (WHO) [4]. Despite these efforts, malaria still causes approximately 600,000 deaths annually, and 95% of these deaths occur in the WHO Africa Region [4].

Vector control policies are typically set by national malaria programs in response to a variety of factors such as local epidemiology and ecology, as well as cost and funding [5–7]. These policies determine where interventions are implemented and the specific interventions used, including the types (e.g. LLIN, IRS, larviciding, etc.), formulations, and distribution methods [8, 9]. This situation forms a heterogeneous landscape of interventions, each of which may impact malaria vectors in different ways in terms of reductions in population size and selection pressures from insecticides that could lead to resistance. Furthermore, the coverage of vector control interventions varies between and within countries [10], exacerbating the potential for differential impacts on local vector populations. The consequences of this heterogeneous vector control depend on the extent to which populations in locations with different interventions or coverage are connected by gene flow, as the rate of gene flow would determine the potential for the impacts of interventions on local vector populations to spread beyond the targeted area and indirectly affect vector populations in other locations.

One approach for understanding gene flow between populations is to infer migration rates based on the genetic differentiation and geographic distance between the populations. Under a model of isolation-by-distance, genetic differentiation would be expected to increase with geographic distance [11] at a rate determined by the dispersal ability of individuals, which is expected to be constant across a homogeneous landscape [12]. However, dispersal ability across a heterogeneous landscape may vary considerably, resulting in populations that are genetically differentiated either more or less than would be expected under isolation-by-distance [12]. Recently developed methods to estimate these deviations from exact isolation-by-distance provide a geospatial visualization of potential corridors and barriers to migration [13, 14]. Using these methods to identify geographical regions with higher or lower migration rates between malaria vector populations could enable predictions of how vector control implemented in one location might indirectly affect vector populations in other locations. Specific examples could include the spread of adaptive alleles conferring insecticide resistance [15], source-sink dynamics [16, 17], or the distribution of future interventions based on genetic modification [18].

The dominant malaria vectors in Africa include *Anopheles gambiae*, *Anopheles coluzzii*, and *Anopheles arabiensis*, which are closely related species in the *A. gambiae* species complex [19]. All three species have large geographical ranges distributed across dozens of countries [20], and in each country the national malaria program independently determines vector control policies. Previous studies of population structure in these *Anopheles* species based on microsatellite loci have generally found relatively low genetic differentiation across thousands of kilometers, indicating less structured populations and high migration rates between locations [21]. However, regions with presumed barriers to migration between populations have also been identified, such as the Rift Valley Complex for *A. gambiae* [22], the Congo Basin tropical rainforest for *A. coluzzii* [23], and the Indian Ocean for *A. arabiensis* [24]. These findings indicate that migration rates between populations of malaria vectors are not uniform across their geographical ranges, but instead depend on the ecological context. Recently, the *Anopheles gambiae* 1000 Genomes (Ag1000G) Project produced whole-genome sequences (WGS) from samples of these species collected in several countries (https://www.malariagen.net/project/ag1000g/). Results of population structure analyses based on single nucleotide polymorphisms (SNPs) from the first [25] and second [26] phases of Ag1000G have provided further evidence that the migration rates in these species depend on the ecological context. Additionally, a comparison between *A. gambiae* and *A. coluzzii* suggests that species-specific ecological adaptations may result in differences in migration rates between species in the same region [26].

In the current study, we assessed population structure and estimated effective migration rates for *A. gambiae*, *A. coluzzii*, and *A. arabiensis* based on WGS data from the third phase of Ag1000G combined with additional, recently processed WGS data. We used principal components analysis (PCA) [26] and ADMIXTURE [27] to assess genetic differentiation between populations based on genome-wide SNPs for each of the three species. We then used the SNP data sets and the mosquito sampling locations to create migration surface maps using the Estimated Effective Migration Surfaces (EEMS) program [13] to visualize where the genetic differentiation between populations within each species was higher or lower than expected under uniform isolation-by-distance. In addition to these assessments of within-species variation in migration rates, we also compared EEMS outputs between species *A. gambiae* and *A. coluzzii* in West and Central Africa, and between *A. gambiae* and *A. arabiensis* in East Africa.

## Results

### Summary of whole-genome sequence data sets

We used the publicly available set of WGS data produced by the third phase of the Ag1000G Project, combined with WGS data from 170 additional *A. gambiae* from Democratic Republic of Congo (DRC) and Malawi, to assess population structure and infer migration rates for three species of malaria vectors in Africa. All sequencing was done on Illumina technology, and sequence reads for all specimens were aligned to the AgamP4 reference genome. Following quality assessment and quality filtration steps (https://malariagen.github.io/vector-data/ag3/ag3.0.html), the final Ag1000G phase 3 data set included 2,784 mosquitoes collected from natural populations in 19 countries. These specimens were assigned to a species (e.g., *A. gambiae*, *A. coluzzii*, or *A. arabiensis*) based on a PCA described in more detail at https://malariagen.github.io/vector-data/ag3/methods.html. We performed analyses separately for each of the three species, excluding specimens assigned to a cryptic taxon (n = 434) and those that appeared to be F1 hybrids (n = 5). We also excluded 75 *A. gambiae*, 21 *A. coluzzii*, and 2 *A. arabiensis* from sibling and parent-offspring kin-groups based on pairwise kinship coefficients using KING version 2.2.7 [27], because these highly-related individuals would otherwise artificially deflate measures of within population genetic differentiation. Our final data set for analysis included 1,565 *A. gambiae* from 107 unique geolocations in 17 countries (**Fig. 1**), 486 *A. coluzzii* (34 geolocations, 9 countries), and 359 *A. arabiensis* (8 geolocations, 4 countries). Some geolocations included samples from more than one species.

**Fig. 1.**
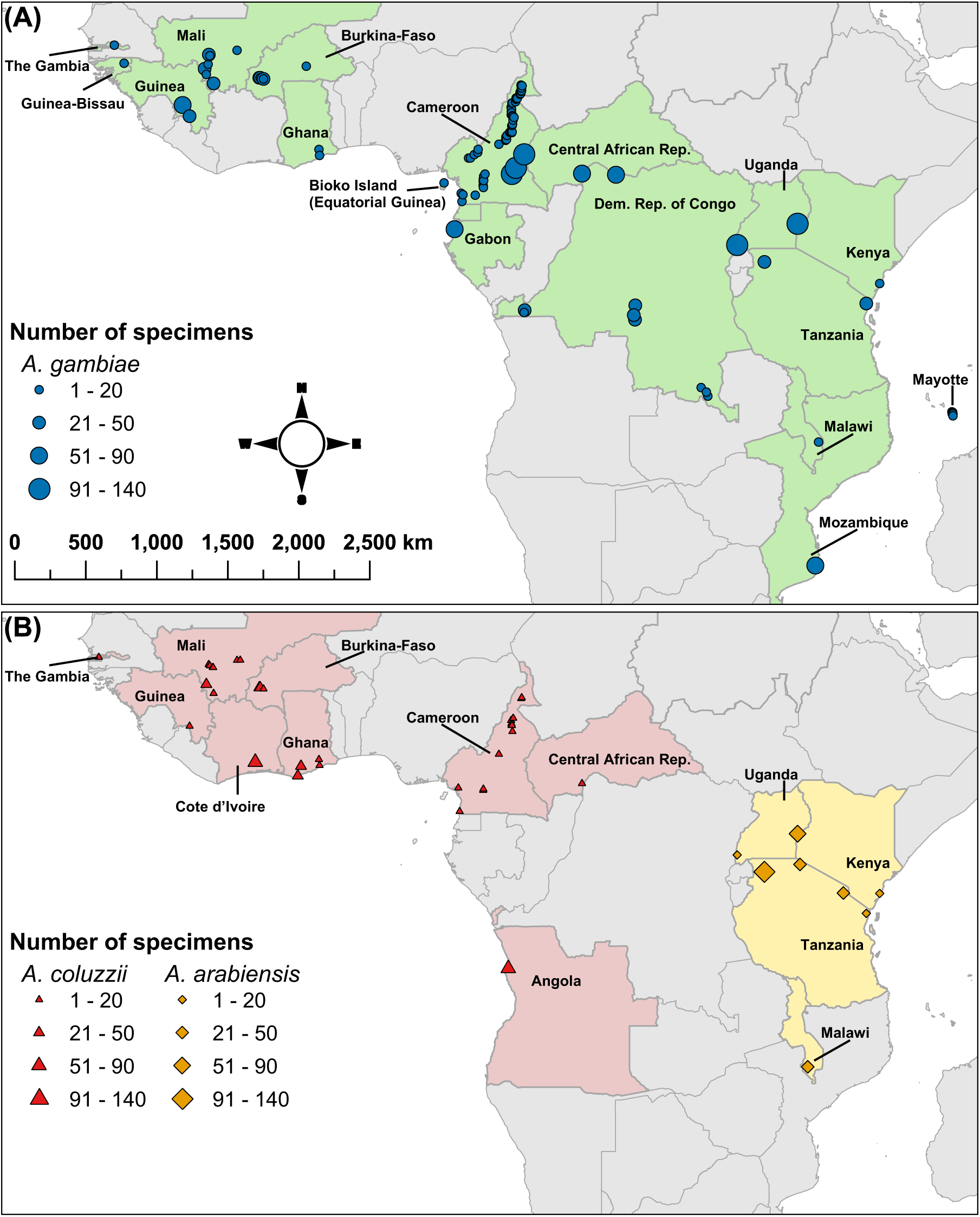
Sample collection locations for (A) *A. gambiae* and (B) *A. coluzzii* and *A. arabiensis*. Symbol shape and color indicate species. Symbol size indicates the number of mosquitoes collected at the location. Green shading (A) indicates countries where *A. gambiae* were collected. Red and yellow shading (B) indicate countries where *A. coluzzii* and *A. arabiensis* were collected, respectively.

The Ag1000G data set includes genome accessibility filters that were developed by applying a machine learning model to colony crosses (https://malariagen.github.io/vector-data/ag3/methods.html). We applied these filters to exclude genomic sites where SNP calling and genotyping was less reliable. The same filter was used for the *A. gambiae* and *A. coluzzii* specimens, while a separate filter was used for *A. arabiensis*. We also excluded genomic regions with polymorphic chromosomal inversions and heterochromatic regions, as these regions of reduced recombination mask signals of geographically associated population structure in these species [25]. Within each species-level data set, we included biallelic positions with a minor allele frequency ≥ 1%, and we removed SNPs in linkage disequilibrium based on an R^2^ threshold of 0.1 in moving windows of 1 kb and a step size of 1 bp. Our final data sets were based on 1,136,140 SNPs in *A. gambiae*, 1,238,862 SNPs in *A. coluzzii*, and 240,091 SNPs in *A. arabiensis*.

### Genome-wide SNP frequencies reveal lower genetic differentiation between *A. gambiae* populations in West Africa and part of Central Africa than the rest of the species geographic range

We assessed population structure within this data set of *A. gambiae* WGS using PCA [28], which revealed at least nine distinct groups along the first three principal components (**Fig. 2A-B**). The first group consisted of samples from West Africa (The Gambia, Guinea-Bissau, Guinea, Mali, Burkina Faso, and Ghana) and northern Central Africa (Cameroon, Central African Republic, and the DRC province of Nord-Ubangi). *A. gambiae* from these locations showed minimal evidence of spatially associated population structure within this large region. Additional clearly distinct groups were formed by samples from 1) Gabon, 2) Kongo-Central Province in southwestern DRC, 3) Kasaï-Central and Haut-Katanga Provinces in southern DRC, 4) Uganda and western Tanzania, 5) coastal Tanzania, coastal Kenya, and Malawi, 6) Mozambique, and 7) Mayotte. *A. gambiae* from these seven regions clustered separately from each other, and from the West/Central Africa group, along the first three principal components. These eight groups were also apparent from the same data set using likelihood-based ADMIXTURE [29] models (**Fig. 2C**), which were performed using values of K (*a priori* clusters) from 3 to 11 (**Fig. S1**). The ninth group apparent from our analysis consisted of samples from Bioko Island, Equatorial Guinea. Although these samples clustered with the West/Central Africa group in PCA along the first six principal components, they were separated as a distinct group along PC8 (**Fig. S2**) and as a mostly distinct group (together with some samples from Cameroon) in the ADMIXTURE results with K≥7 (**Fig. 2C, Fig. S1**).

**Fig. 2.**
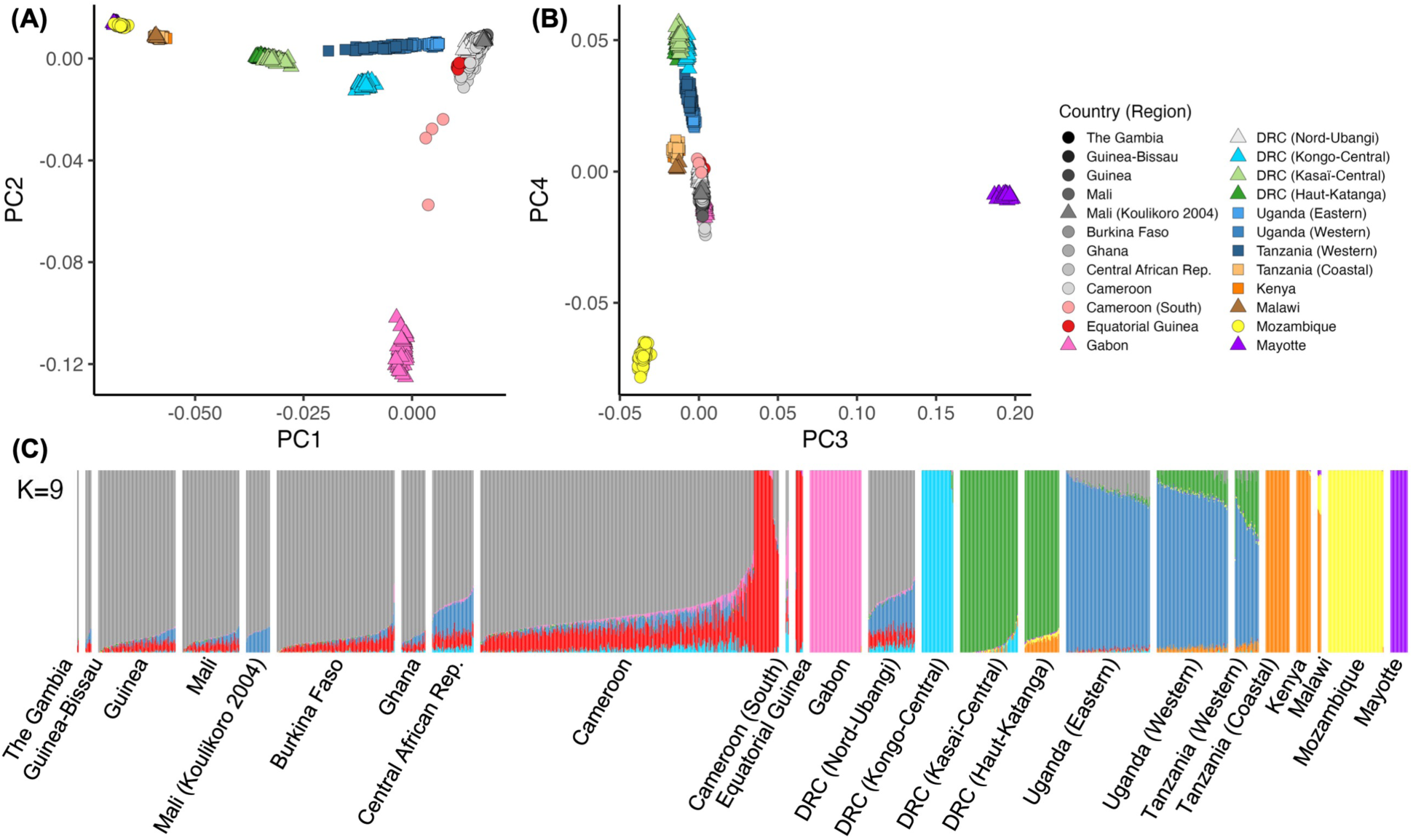
Population structure of *A. gambiae*. (**A**) Principal components (PC) 1 and 2. (**B**) PCs 3 and 4. Each point within a PCA panel represents a single mosquito. Symbol color and shape indicates the country (or region within a country) where the sample was collected. (**C**) ADMIXTURE with K=9. Each bar represents an individual mosquito, with bars grouped by the country (or region within a country) where the mosquitoes were collected.

The PCA and ADMIXTURE results also showed more subtle genetic differentiation within the groups described above, suggesting population sub-structure within these groups of *A. gambiae*. First, we observed two sub-groups within the Uganda and western Tanzania group. In the ADMIXTURE results, one ancestral population contributed 43-100% of the modeled ancestry for *A. gambiae* within the Uganda and western Tanzania group (**Fig. 2C**). In terms of sub-structure, *A. gambiae* from eastern Uganda showed shared ancestry with the West/Central Africa group, which includes Nord-Ubangi Province in northern DRC, while *A. gambiae* from western Uganda and western Tanzania showed shared ancestry with Kasaï-Central and Haut-Katanga Provinces in southern DRC. These results were consistent with the relative positions of these samples along PC1 in the PCA results (**Fig. 2A**).

We observed additional sub-structure in East Africa between *A. gambiae* from coastal Tanzania and Kenya and those from Malawi. In the ADMIXTURE results, one ancestral population contributed 97-100% of the modeled ancestry for *A. gambiae* from coastal Tanzania and Kenya, while *A. gambiae* from Malawi showed shared ancestry (19-22%) with those from Mozambique (**Fig. 2C**). In the PCA results, *A. gambiae* from Malawi separated as a distinct group along PC6 (**Fig. S2**).

Within Cameroon the strongest evidence for population sub-structure was between *A. gambiae* from South Region and those from the remaining administrative regions of the country, although our interpretation is limited by the number of specimens from South Region (n = 4). In the PCA results, these individuals from South Region appeared between *A. gambiae* from the remaining locations in Cameroon and those from Gabon along PC2 (**Fig. 2A**), and in the ADMIXURE results the individuals from South Region appeared to share an equal amount of ancestry between those same two groups (**Fig. 2C**). While we also observed some additional evidence of population structure within Cameroon, e.g., along PC5 and PC10 (**Fig. S2**) and with ADMIXTURE from K≥7 (**Fig. S1**), these differentiated specimens were not clustered geographically.

Within the West/Central Africa group, the ADMIXTURE model indicated that most *A. gambiae* shared common ancestry with the Uganda and western Tanzania group for a small proportion of their genome (**Fig. 2C**). Notably, there was an east-to-west gradient in the proportion of ancestry attributed to this group, with samples from Guinea, Mali, Burkina Faso and Ghana showing lower proportions than those from Central African Republic and Nord-Ubangi Province in northern DRC. Additionally, *A. gambiae* collected in 2004 from Koulikoro Region, Mali (n = 32) showed subtle, but still mostly distinct, genetic differentiation from the remaining *A. gambiae* collected in West Africa according to ADMIXTURE (**Fig. 2C** and **Fig. S1**) as well as PCA along PC9 (**Fig. S2**). Nearby collections included specimens from Segou Region, Mali (n = 1) and Centre-Sud Region, Burkina Faso (n = 13) in 2004, and specimens from Mali, Guinea and Burkina Faso in 2012 and 2014, including 33 specimens from Koulikoro Region, Mali, all of which were more similar to each other than the Koulikoro 2004 specimens. Potential explanations for these subtle differences include genetic changes over time in *A. gambiae* populations in Koulikoro, or long-distance migration [30] from an unsampled location to Koulikoro. However, the available data are not sufficient to distinguish between these or other hypotheses.

### Migration rates between *A. gambiae* populations vary across the species geographic range

We used EEMS to create maps of effective migration rates for *A. gambiae*, thereby visualizing where the genetic differentiation between the populations of *A. gambiae* described above was higher or lower than expected based on the geographic distance between the sampling locations. The null model in EEMS is that any observed differentiation between populations is the result of isolation-by-distance, and the mean rate of increase in differentiation across all populations determines the mean migration rate across the geographic range being mapped. The effective migration rates (*m*) modeled by EEMS are parameterized as deviations from this mean rate (*µ*), so that log_10_(*m*) = 0 is equivalent to the overall mean rate, and log_10_(*m*) = 1, for example, corresponds to effective migration at a rate ten times higher than the overall mean [13]. We found that the genetic differentiation among *A. gambiae* from locations in the West/Central Africa group described above was generally much less than would be expected under isolation-by-distance (**Fig. 3**). Consequently, the effective migration rates between locations in this region estimated by EEMS were generally higher than, or at least equal to, the mean rate across all sampled locations. In addition to the map visualizing the point estimates of log_10_(*m*) across space (**Fig. 3A**), the EEMS model also maps exceedance probabilities, showing where the posterior probability of the log_10_(*m*) being greater than or less than 0 exceeds 0.90 or 0.95 (**Fig. 3B**). This latter map provides a visualization of where migration rates are most likely to depart from the overall mean rate of log_10_(*m*) = 0, thereby indicating regions with the highest confidence migration barriers or corridors based on the EEMS model. For example, there are several areas within the West/Central Africa group of *A. gambiae* where the probability of a higher than expected migration rate is greater than 0.90, while there are none with this high probability of a lower than expected migration rate.

**Fig. 3.**
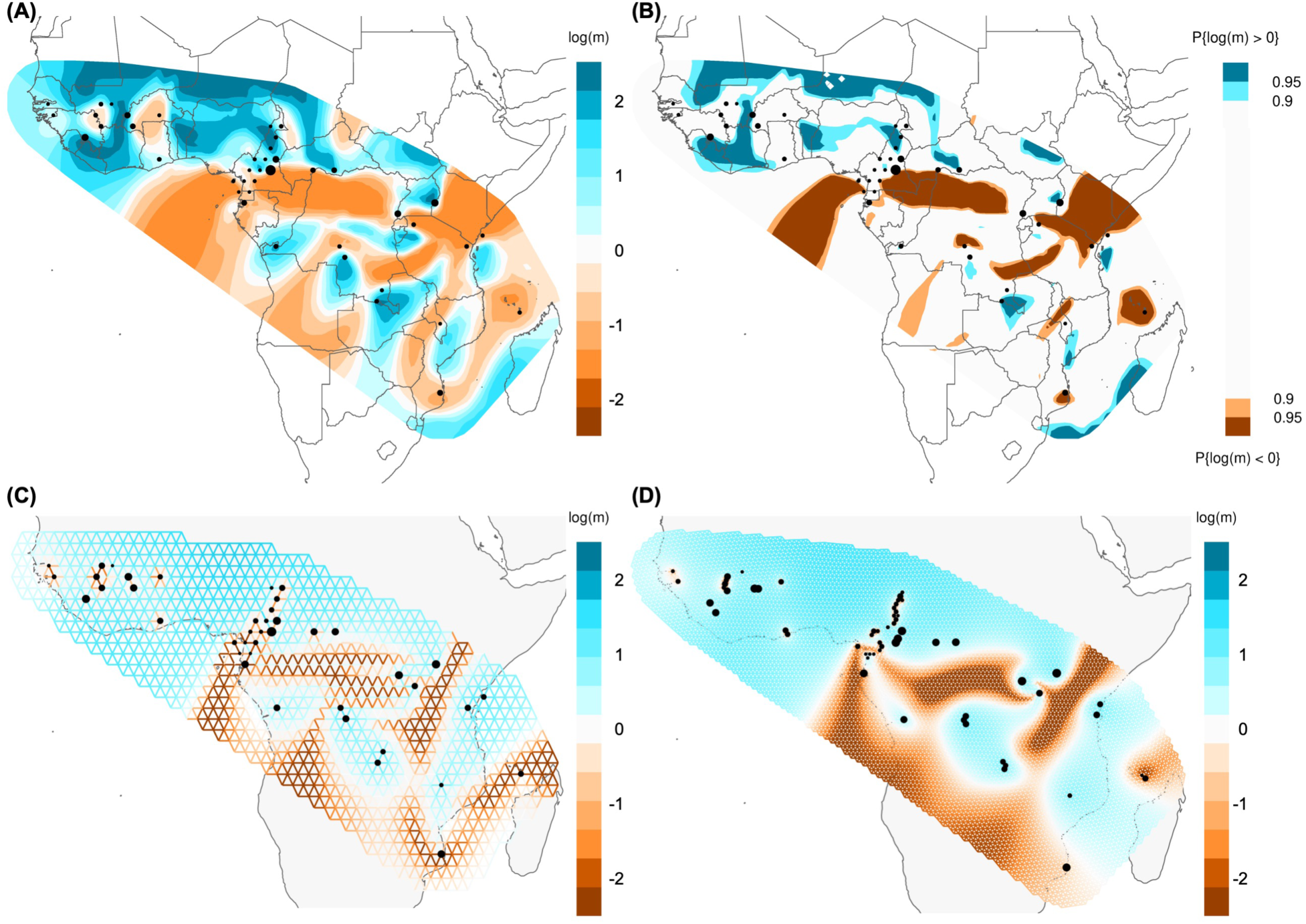
Estimated effective migration surfaces for *A. gambiae*. (**A**) Posterior mean effective migration rates, log10(*m*), from EEMS. (**B**) Posterior probabilities of the effective migration rate, log10(*m*), being greater than or less than 0 from EEMS. (**C**) Effective migration rates, log10(*m*), from FEEMS using the same grid resolution as panel **A**. (**D**) Effective migration rates, log10(*m*), from FEEMS using a higher resolution grid, with edge lengths of 55 km. Black circles show the geolocations of samples included in the analysis, aligned to the nearest node on the grid, with circle size scaled to the number of samples.

Outside of the West/Central Africa group, we found that the genetic differentiation between *A. gambiae* from the other populations described above was generally higher than expected under isolation-by-distance, resulting in lower-than-average effective migration rates estimated by EEMS. These lower-than-average migration rates indicate the presence of barriers to migration and/or absence of facilitators of migration for *A. gambiae* moving between these regions. For example, EEMS estimated lower than average migration rates for *A. gambiae* across the Rift Valley Complex in East Africa, between northern and southern areas of Central Africa, and between Bioko Island and Cameroon. This last example highlights additional insights possible from spatially explicit analyses such as EEMS. While the results from PCA and ADMIXTURE indicated relatively subtle genetic differentiation between *A. gambiae* from Bioko Island and Cameroon, the results from EEMS indicate that those differences were higher than expected under isolation-by-distance, presumably due to the roughly 30 km of ocean acting as a barrier to migration between Bioko Island and the mainland. A second example is that the effective migration rates between the West/Central Africa group and eastern Uganda were likely equal to, or slightly higher than, the mean rate expected under isolation-by-distance. Although the PCA and ADMIXTURE results showed distinct genetic differentiation between *A. gambiae* from the West/Central Africa group and those from eastern Uganda, the results from EEMS (**Fig. 3A-B**) indicated that these differences may be due to isolation-by-distance.

### Migration rate estimates were robust to changes in grid size and jackknife resampling

We investigated the robustness of our estimated effective migration rate maps in several ways using Fast Estimation of Effective Migration Surfaces (FEEMS), a method that is conceptually similar to EEMS but which uses a penalized likelihood framework to allow more efficient optimization and output [14]. One limitation of EEMS is that it is computationally intensive, whereas the higher computational efficiency of FEEMS allows for estimation of effective migration rate maps at high throughput and with much finer geospatial resolution. In both EEMS and FEEMS, estimates of migration rates are made along a triangular grid, where grid vertices represent demes in a stepping-stone model, and each sampling location is assigned to the nearest vertex on the grid.

First, we compared the outputs of EEMS and FEEMS by running both models using the same 1000-deme grids, which is the finest resolution possible with EEMS. The effective migration surfaces estimated by EEMS (**Fig. 3A**) and FEEMS (**Fig. 3C**) for *An. gambiae* were generally similar, especially in areas nearest to the geolocations where samples were collected. Both methods estimated lower than average migration rates for *A. gambiae* across the Great Rift Valley in East Africa, between northern and southern areas of Central Africa, and between Bioko Island and Cameroon; as well as higher than average migration rates across most of West Africa. Although there were also areas where the estimates of effective migration for *A. gambiae* differed between the two methods, these differences generally occurred where the exceedance probability map from EEMS (**Fig. 3B**) indicated lower confidence that the estimates were different from 0 (i.e., P(log(*m*) ≠ 0) < 0.90).

To assess the effect of grid resolution on model outcomes [31], with more vertices (demes) than possible in EEMS, we ran FEEMS using a range of grid resolutions, with edge lengths of 55, 110 and 220 km for each species. The 220 km edges corresponded with the finest resolution possible in EEMS, while each halving of the edge length increased the number of grid nodes by approximately four-fold. The results from FEEMS were robust to changes in the grid resolution when comparing across edge sizes from 55 km to 220 km (**Fig. 3C-D, Fig. S3-S5**), with both high and low migration estimates occurring in the same regions across all three grids. Notably, the FEEMS results were more dependent on the smoothing parameter, lambda, than on the grid resolution (**Fig. S3-S5**). In the main text (**Fig. 3C-D**), we have presented results from FEEMS using the value for that had the lowest mean square cross-validation error across a range of lambda from 0.1 to 100 (**Fig. S3-S5**).

Finally, we visually assessed the resulting maps from each iteration of jackknife cross-validation to assess the robustness of FEEMS estimates to removal of observed specimens from a single deme (**Supplementary file Ag FEEMS jackknife maps**). In general, removing a single observed deme from the FEEMS model had a greater influence on the surrounding migration rate estimates for demes with fewer nearby neighboring observed demes. In other words, migration rates estimated by FEEMS were generally robust to the removal of data from a single location when data were available for other nearby locations. One exception to this result was the influence of the *A. gambiae* specimens collected near Koulikoro, Mali, in 2004. In this case, the genetic differentiation between these specimens and those from several nearby locations caused FEEMS to estimate relatively low migration rates near Koulikoro. When the Koulikoro specimens were removed, the FEEMS model estimated relatively high migration rates in this region. As with the PCA and ADMIXTURE results, potential explanations for the different estimates of effective migration rates here include genetic changes over time in *A. gambiae* populations in Koulikoro, or long-distance migration [30] from an unsampled location to Koulikoro.

### *Anopheles coluzzii* in West and Central Africa had higher genetic differentiation between populations than *A. gambiae* in the same region

The spatial extent of locations where the *A. coluzzii* were collected included most of the West/Central Africa region where *A. gambiae* showed very little spatially associated population structure. Over that same region, we found distinct genetic differentiation between *A. coluzzii* from different locations according to the results of PCA (**Fig. 4A**) and ADMIXTURE (**Fig. 4B**). In some cases, e.g. between coastal and inland populations, those differences were greater than would be expected under isolation-by-distance, according to our analysis using EEMS (**Fig. 5**).

**Fig. 4.**
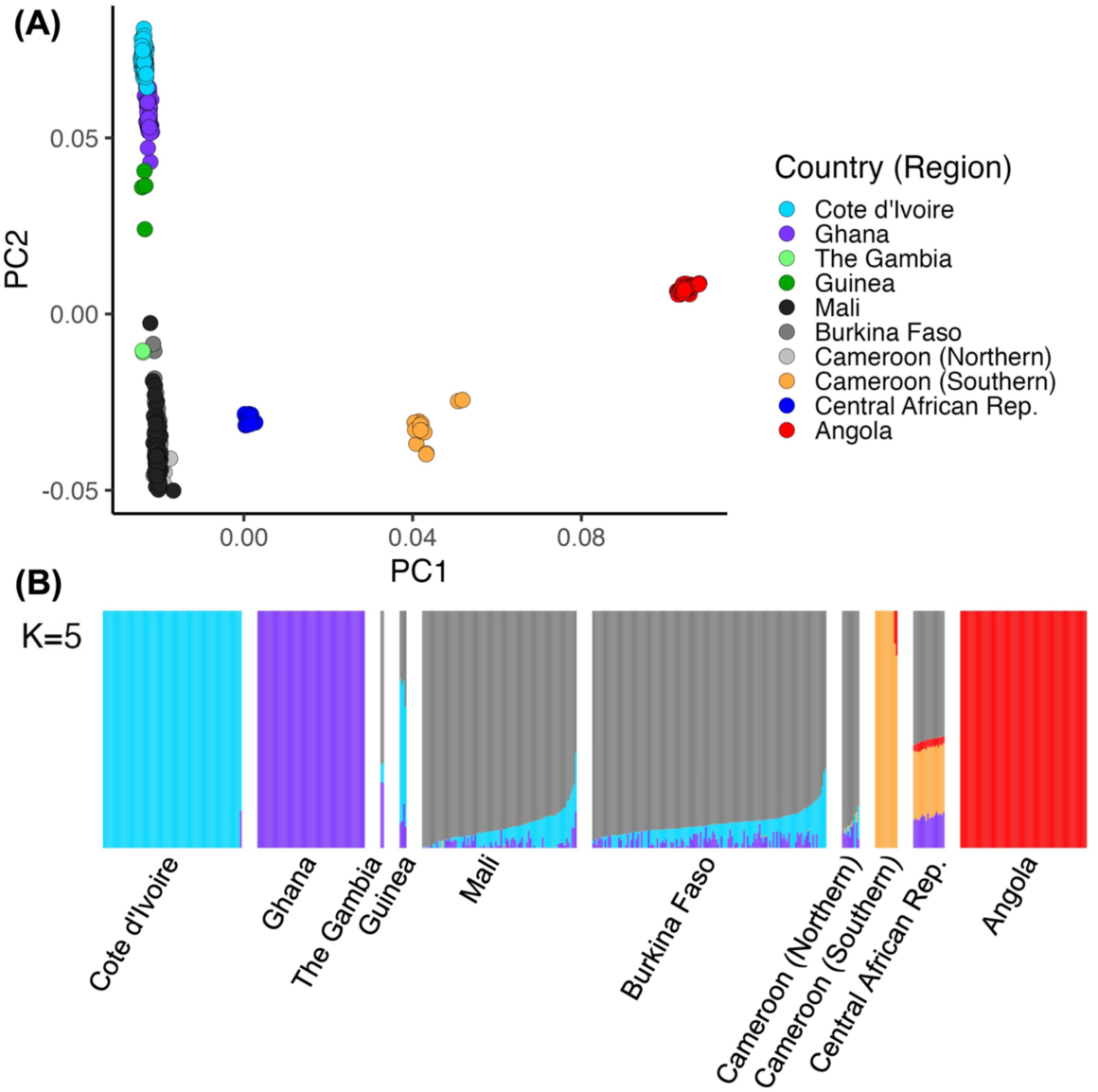
Population structure of *A. coluzzii*. (**A**) Principal components (PC) 1 and 2. Each point represents a single mosquito. Symbol color indicates the country (or region within a country) where the sample was collected. (**B**) ADMIXTURE with K=5. Each bar represents an individual mosquito, with bars grouped by the country (or region within a country) where the mosquitoes were collected.

**Fig. 5.**
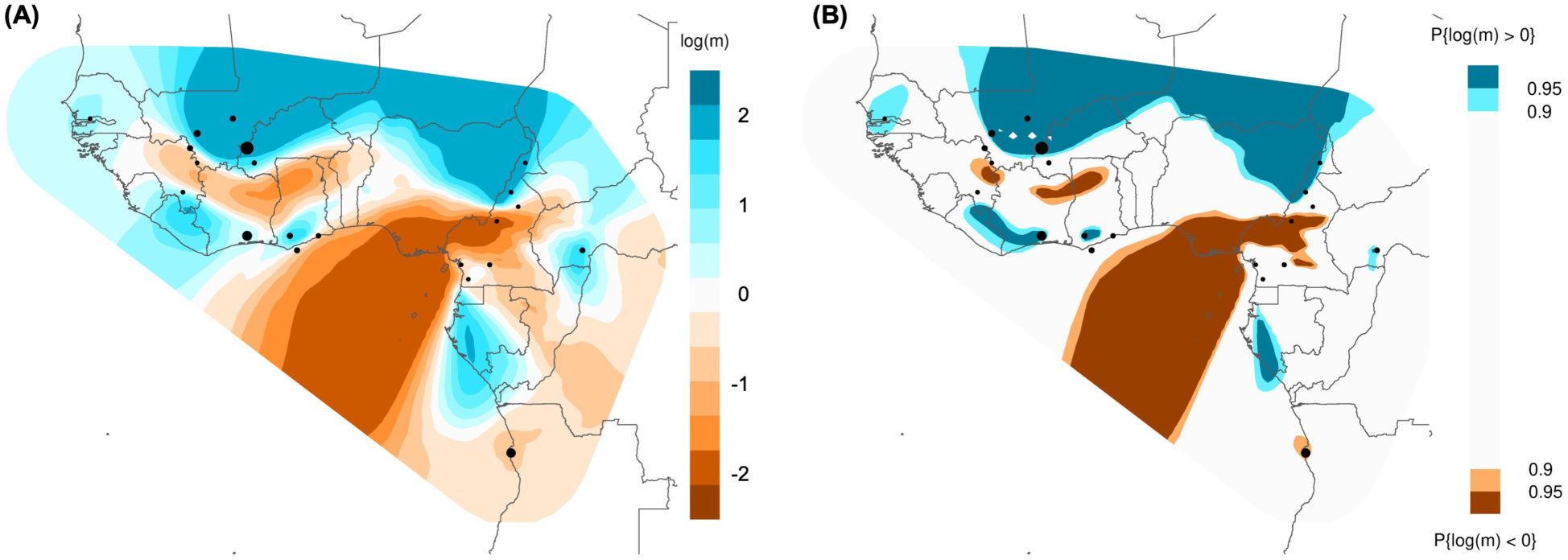
Estimated effective migration surfaces for *A. coluzzii*. (**A**) Posterior mean effective migration rates, log_10_(*m*), from EEMS. (**B**) Posterior probabilities of the effective migration rate, log_10_(*m*), being greater than or less than 0 from EEMS. Black circles show the geolocations of samples included in the analysis, aligned to the nearest node on the grid, with circle size scaled to the number of samples.

Four broader geographic groups of *A. coluzzii* were apparent: 1) locations nearer to the coast of West Africa in Ghana and Côte d’Ivoire; 2) locations nearer to the Sahel region of West and Central Africa, including Mali, Burkina Faso and three northern regions of Cameroon (Far North, North, and Adamawa); 3) three southern regions of Cameroon (Centre, Littoral, and South); and 4) Angola. In addition to these broader groups, *A. coluzzii* from Guinea and Central African Republic provide two examples of populations that may be admixtures of separate ancestral populations. First, *A. coluzzii* from Guinea clustered between the coastal West Africa and Sahelian individuals in PCA (**Fig. 4A**), while the ADMIXTURE results showed *A. coluzzii* from Guinea sharing genetic patterns of both regions for K values from 3 to 8 (**Fig. 4B** and **Fig. S6**). Second, *A. coluzzii* from Central African Republic formed a distinct cluster in the PCA plots, clustering between individuals from southern Cameroon and individuals from northern Cameroon along PC1 (**Fig. 4A**), and they showed ancestry matching both southern and northern Cameroon in the ADMIXTURE results for values of K from 2 to 5 (**Fig. 4B** and **Fig. S6**).

We found the highest estimated effective migration rates among *A. coluzzii* from locations near the Sahel (Mali, Burkina Faso and northern Cameroon), based on results from EEMS (**Fig. 5**) and FEEMS (**Fig. S7-S10**). For *A. coluzzii* from Ghana and Côte d’Ivoire, the effective migration rates between these geolocations estimated by EEMS were higher than average, indicating that the subtle genetic differences between samples from these two countries, observed in PCA and ADMIXTURE, were less than expected given the geographic distance. The lowest estimated effective migration rates were observed between the geographic groupings outlined above. Notably, these estimates were relatively close to the mean migration rate in some cases, for example between *A. coluzzii* from southern Cameroon and Angola (**Fig. 5**). Given the long distance between those two locations, this indicates that the clear genetic differentiation between the respective populations is approximately as much as would be expected under uniform isolation-by-distance.

### *Anopheles arabiensis* in East Africa had lower genetic differentiation between populations than *A. gambiae* in the same region

The data set analyzed here included fewer specimens of *A. arabiensis* (n = 359) from fewer unique sampling locations (eight) than the other two species, and the locations covered only a portion of the known species range of *A. arabiensis*. Still, the sampling locations mostly overlapped with those of *A. gambiae* in East Africa, providing an opportunity for comparisons between these two species. We observed distinct genetic differentiation between *A. arabiensis* from three geographic groupings: 1) Malawi; 2) eastern Uganda; and 3) Kenya and Tanzania (**Fig. 6**). Additionally, the geolocation in western Uganda, which only had one specimen of *A. arabiensis*, grouped with *A. arabiensis* from Kenya and Tanzania. The results from EEMS (**Fig. 7A**) and FEEMS (**Fig. S12-S15**) suggest similar groupings, with *A. arabiensis* from Malawi and eastern Uganda separated from the other regions by areas of low migration rates. However, the exceedance probability map from EEMS (**Fig. 7B**) shows less than 90% probability across most of the study area that the migration rates differ from the mean rate expected by isolation-by-distance, possibly due to the low number of specimens and locations in this data set.

**Fig. 6.**
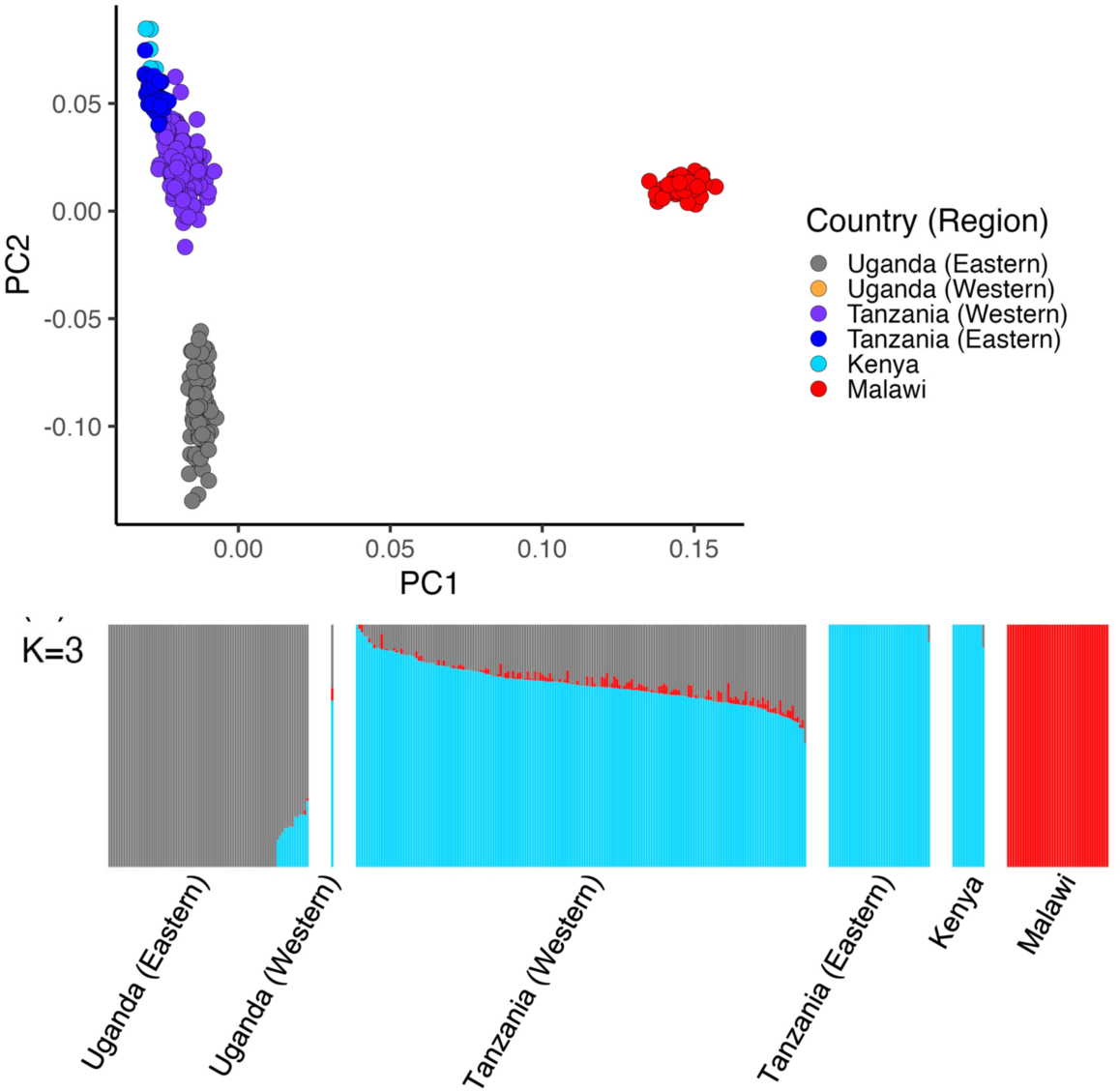
Population structure of *A. arabiensis*. (**A**) Principal components (PC) 1 and 2. Each point represents a single mosquito. Symbol color indicates the country (or region within a country) where the sample was collected. (**B**) ADMIXTURE with K=5. Each bar represents an individual mosquito, with bars grouped by the country (or region within a country) where the mosquitoes were collected.

**Fig. 7.**
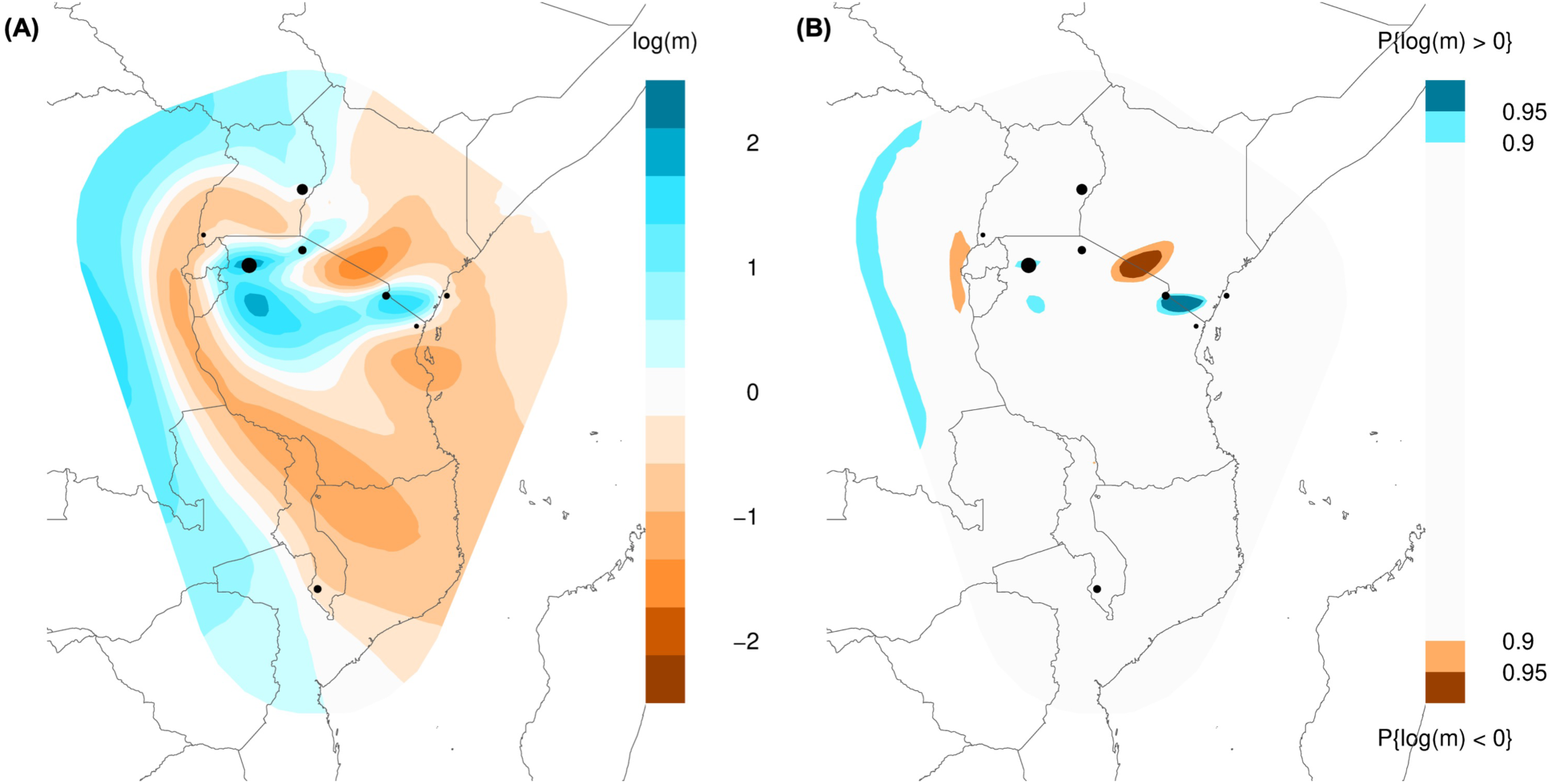
Estimated effective migration surfaces for *A. arabiensis*. (**A**) Posterior mean effective migration rates, log_10_(*m*), from EEMS. (**B**) Posterior probabilities of the effective migration rate, log_10_(*m*), being greater than or less than 0 from EEMS. Black circles show the geolocations of samples included in the analysis, aligned to the nearest node on the grid, with circle size scaled to the number of samples.

Within the Kenya and Tanzania group, there was some evidence of additional sub-structure, but it was relatively subtle. *A. arabiensis* from western Tanzania (Muleba/Kagera and Tarime/Mara) formed a cluster along PC2, which mostly does not overlap with *A. arabiensis* from eastern Tanzania (Moshi/Manyara and Muheza/Tanga) and Kenya (Kilifi). The ADMIXTURE results show a similarly subtle difference between *A. arabiensis* from these regions (**Fig. 6B** and **Fig. S11**). This same pattern is apparent in the results from EEMS, with an area of lower effective migration estimated in northern Tanzania between samples from Tarime (to the west) and Moshi (to the east; **Fig. 7**). However, according to the FEEMS results (**Fig. S12-S15**), the genetic differentiation between samples from Moshi in eastern Tanzania and Muleba and Tarime in western Tanzania was less than expected under isolation-by-distance (i.e., high estimated effective migration), while the genetic differentiation between samples from Moshi and coastal Tanzania and Kenya were greater than expected based on the geographic distance (i.e., low estimated effective migration).

## Discussion

In this study, we assessed the population structure of three African malaria vector species and estimated effective migration rates between populations within each species. We found that the genetic differentiation observed between populations of *A. gambiae* from East, Southern and part of Central Africa was greater than would be expected under uniform isolation-by-distance, with relatively low effective migration rates estimated between populations in these regions. These results contrast with our observations of *A. gambiae* populations in West and northern Central Africa, which had low genetic differentiation despite the long geographic distances between them, corresponding with relatively high effective migration rates in these regions. In contrast to *A. gambiae*, we found greater genetic differentiation between *A. coluzzii* populations in West and Central Africa, indicating that these closely related species respond to, or interact with, barriers and/or facilitators of migration in this region differently. Conversely, we found less distinct population structure between *A. arabiensis* specimens from East Africa, especially within Tanzania, compared to *A. gambiae*. These results demonstrate that migration rates between populations vary across geographic distributions of these important malaria vectors, likely in response to ecological barriers or facilitators of migration. Furthermore, these apparent effects on migration differ between species, even for very closely related species.

This is the first study to use the EEMS framework [13, 14] to elucidate the spatial population structure of a disease vector. While PCA and ADMIXTURE allowed us to visualize the relative genomic similarity of the sampled vectors and thereby identify groups of sampling locations with higher genetic similarity compared to the other groups, PCA and ADMIXTURE do not account for the geographic distances between sample collection sites. By using EEMS to explicitly model both the genomic data and the sampling locations, we were able to visualize how migration rates likely vary across the landscape to produce the observed patterns of relative genetic differentiation among the sampled geolocations. While each of these methods has its own strengths and weaknesses, we have focused here on findings that were consistent across the methods, for which we have the highest confidence. Additionally, we have highlighted findings from EEMS that differ from the PCA and ADMIXTURE results when these were plausibly explained by the geographic distances between sample collection sites.

This study expands on previous assessments of population structure in the *A. gambiae* species complex based on earlier phases of the Ag1000G project [25, 26]. Those studies found low genetic differentiation between *A. gambiae* from West Africa (Guinea, Burkina Faso, and Ghana), Cameroon and Uganda; and they showed greater genetic differentiation between samples from Bioko Island (Equatorial Guinea), Gabon, Mayotte, and Coastal Kenya, and between each of those locations and West/Central Africa. With the addition of *A. gambiae* specimens from Mali, Central African Republic, Democratic Republic of Congo, Tanzania, Malawi, and Mozambique, our updated analyses largely align with the previous findings: *A. gambiae* across West and northern Central Africa show very little genetic differentiation, while samples from locations outside this region show more distinct genetic differentiation. Our results here differ slightly from the previous findings, in that we found distinct genetic differentiation between *A. gambiae* from Uganda and the West/Central Africa group. However, this genetic differentiation was less than we observed between other populations, and our EEMS and FEEMS analyses suggest that migration between populations of *A. gambiae* in Uganda and Central Africa (e.g. northern DRC, Central African Republic and Cameroon) may be occurring at roughly the rate expected under isolation-by-distance, or even slightly higher. Ultimately, denser mosquito sampling is needed to better delineate the population structure of *A. gambiae* across the geographic distribution of this malaria vector species. As the availability of *A. gambiae* genomic sequences from previously unrepresented regions continues to increase, we expect additional geographically associated population structure to be revealed.

Phase 2 of the Ag1000G project [26] showed differentiation between *A. coluzzii* specimens from coastal West Africa (Ghana and Côte d’Ivoire) and further inland (Burkina Faso). With the addition of specimens from Mali and Cameroon in the phase 3 data set, our analyses suggest a broader pattern: *A. coluzzii* nearer to the Sahel region of West and Central Africa (Mali, Burkina Faso and northern Cameroon) show very little genetic differentiation from each other but show distinct genetic differentiation from *A. coluzzii* nearer to the coast (Ghana, Côte d’Ivoire and southern Cameroon). While the high effective migration rates estimated here for *A. coluzzii* from the near-Sahel region are consistent with the estimates for *A. gambiae* in the same region, we found differences between the two species in the way these near-Sahel populations are connected, or not, to coastal populations, with *A. coluzzii* having lower effective migration rates between the coast and inland, and *A. gambiae* being more connected. These differences in population connectivity between two closely related species in the same region suggest that different factors may either impede or facilitate successful migration in each species. Potential factors could include long-distance windborne dispersal [30] or adaptations to local habitats [32]. One limitation of this comparison between species was that the number of specimens, and the number and location of sampling sites, differed between species. Still, there was considerable spatial overlap, with both *A. gambiae* and *A. coluzzii* collected from 28 of the 34 sites with *A. coluzzii*.

This is the first study to look at the population structure of *A. arabiensis* using whole-genome sequences. The Ag1000G project phase 3 data set used in this study includes samples from East and Southern Africa, and our results are broadly consistent with previous studies of *A. arabiensis* population structure in this region based on microsatellites. Donnelly and Townson [33] covered a relatively broad spatial scale, assessing samples of *A. arabiensis* from nine locations in five countries, from Sudan in the north to Mozambique in the south, and found five groupings of sub-populations that each aligned with the country where the samples were collected. These results showed evidence of population structure at a broad spatial scale across the region, consistent with our study. At a finer scale, Maliti *et al.* [34] assessed *A. arabiensis* and *A. gambiae* in central, coastal and island regions of Tanzania, and they found less population structure in *A. arabiensis* in this area than for *A. gambiae*. Similarly, Kamau *et al.* [35] found genetic differentiation between samples of *A. gambiae* in western and coastal Kenya, but not *A. arabiensis*. Our finding of less distinct population structure between samples of *A. arabiensis* from East Africa, compared to *A. gambiae*, is consistent with these previous studies, despite the limited scope of our *A. arabiensis* data set.

Our analyses were based on a large number of biallelic SNPs with a minor allele frequency ≥ 1% (> 1 million SNPs in *A. gambiae* and *A. coluzzii*, and > 200,000 SNPs in *A. arabiensis*). The signal from analyses of population structure from this type of data is likely driven by migration over several generations stretching to the relatively distant past due to reasons including: (1) the relatively old ages of common neutral variants [36], and (2) the use of SNP-based genetic distance metrics that capture mean coalescent times over a long history instead of the frequencies of recent coalescent events [37]. Future analyses using metrics capable of capturing more recent information, such as rare variants with younger ages [38] or longer shared segments of the genome that are identical-by-descent and that reflect more recent recombination and coalescent events [37, 39–42], could provide more insights into recent gene flow.

In conclusion, our results, supported by those of several previous studies, establish that migration rates between malaria vector populations vary both between and within species, presumably based on species-specific responses to ecological or environmental conditions that may impede or facilitate migration, and the geographical patterns of these conditions across the landscape. These differences within and between species have potential implications for malaria research and control programs planning surveillance of vector populations, monitoring for insecticide resistance, and evaluating interventions. These findings should also be considered when planning multi-country malaria control strategies, e.g. the Elimination Eight initiative [43], or developing novel control strategies that rely on mosquito mating, e.g. genetic modification [18]. These findings also raise questions about which environmental features may be acting as migration barriers or corridors for malaria vectors, and whether these features may change over time.

## Methods

### Whole genome sequencing and SNP calling

Whole genomes of 4,693 individual mosquitoes were sequenced at the Wellcome Sanger Institute for the Ag1000G phase 3 data set. Sequencing was done on the Illumina HiSeq 2000 platform (100 bp paired-end reads) or Illumina HiSeq X platform (150 bp paired-end reads). Sequence reads were aligned to the AgamP4 reference genome using BWA version 0.7.15. Indel realignment and SNP calling were done using GATK version 3.7-0. Mosquitoes were excluded if median coverage across all genome positions (across all chromosomes) was less than 10x, or if less than 50% of the reference genome was covered by at least 1x. Additional mosquitoes were excluded based on analyses to identify cross-contamination and population outliers. More details about the Ag1000G project, including descriptions of the partner studies and quality assessment and quality filtration steps for the whole-genome sequence data are available at https://malariagen.github.io/vector-data/ag3/ag3.0.html. The final Ag1000G phase 3 data set included 2,784 mosquitoes collected from natural populations in 19 countries.

We also included 165 additional *A. gambiae* from 8 locations in Democratic Republic of Congo and 5 additional *A. gambiae* from 1 location in Malawi. The 165 specimens from Democratic Republic of Congo were sequenced at Wellcome Sanger Institute on an Illumina HiSeq (150 bp paired-end reads) as part of the Vector Observatory project under sample set ID 1264-VO-CD-WATSENGA-VMF00164 as described in Dennis *et al*. [44]. The 5 *A. gambiae* from Malawi were sequenced at the University of Marland on the Illumina NovaSeq 6000 platform (150 bp paired-end reads). Sequence alignment, indel realignment, SNP calling, and all quality assessment and filtration steps for these 170 specimens were performed following the same steps as used for the Ag1000G phase 3 data set.

### Included samples and genomic positions

Analyses described here were done separately for each species. Specimens were assigned to each species according to a principal components analysis (PCA) described in more detail at https://malariagen.github.io/vector-data/ag3/methods.html. Specimens assigned to a cryptic taxon and those that appeared to be F1 hybrids were excluded from this study.

Genome accessibility filters were developed by applying a machine learning model to colony crosses (https://malariagen.github.io/vector-data/ag3/methods.html), and these filters were applied to exclude genomic sites where SNP calling and genotyping was less reliable. The same filter was used for *A. gambiae* and *A. coluzzii*, while a separate filter was used for *A. arabiensis*. Pairwise kinship coefficients were estimated using KING version 2.2.7 to identify first-degree relatives (full siblings or parent-offspring). These kinship analyses were based on the euchromatic regions of the autosomal chromosomes, using only biallelic positions with a minor allele frequency ≥ 1%. After identifying all first-degree relative pairs, kin-groups were constructed such that each group contained all first-degree relatives from the data set for all individuals in the kin-group. All but one mosquito from each kin-group were excluded from further analysis, leaving 1,565 *A. gambiae*, 486 *A. coluzzii*, and 359 *A. arabiensis*.

For all subsequent analyses, we excluded genomic regions with polymorphic chromosomal inversions in addition to excluding heterochromatic regions. For *A. gambiae* and *A. coluzzii*, we used genomic positions on chromosome 3R from 1-37 Mb and on chromosome 3L from 15-41 Mb. Due to an additional inversion polymorphism on chromosome 3R in *A. arabiensis*, we only included chromosome 3L (15-41 Mb) in that species. Within each species-level data set, we only included biallelic positions with a minor allele frequency ≥ 1%, and we removed SNPs in linkage disequilibrium based on an R^2^ threshold of 0.1 in moving windows of 1 kb and a step size of 1 bp.

### Spatial data

The geolocation for each sampled mosquito was recorded as part of each contributing study to the Ag1000G project (https://malariagen.github.io/vector-data/studies-ag1000g.html). Sampling locations for the 165 *A. gambiae* collected in Democratic Republic of Congo (sample set ID 1264-VO-CD-WATSENGA-VMF00164) are described by Dennis *et al.* [44]. The five *A. gambiae* from Malawi were collected as blood-fed adult females from inside houses near Ntaja Health Center in Machinga District, Malawi, in March-April 2022. In most contributing studies, all mosquitoes sampled from the same village were assigned the same GPS coordinates, i.e., for that village, such that the coordinates are accurate to within a few hundred meters, at worst. For 5 out of 128 unique geolocations, all mosquitoes from the same county or district were assigned to the same GPS coordinates, such that those coordinates are only accurate to within ∼20-50 km. Notably, for those 5 sample geolocations, the next nearest neighbor was ∼200-800 km away, and therefore the GPS coordinates were sufficiently accurate for our analyses.

For generation of estimated effective migration surfaces (EEMS and FEEMS), the geographic boundary of each map was defined as a convex hull 600 km beyond the outermost geolocations in each sample set to reduce potential edge effects.

### Population structure

PCA was performed using PLINK version 1.90b6.18. Within each species, we plotted the principal components with the highest eigenvalues (explaining the highest percentage of the variance) after examining the scree plot. For all PCA plots, individual samples were color-coded by country (or administrative regions when applicable) to visually assess potential associations between genetic clustering and sample geolocations.

Admixture models were performed using ADMIXTURE version 1.3.0 for values of K (clusters) from 3 to 11 (*A. gambiae*), 2 to 8 (*A. coluzzii*) and 1 to 5 (*A. arabiensis*), with the range of K based on assessment of PCA results. For each value of K within each species, we ran ten replicates of ADMIXTURE, each with a different random seed, and conducted 5-fold cross validation on each replicate. We set the point estimation algorithm to stop when the log-likelihood increased by less than ε = 10^-4^ between iterations (the default). Within each value of K, we plotted the run with the highest likelihood, grouping samples by country (or administrative regions where applicable) to visually assess potential associations between genetic clustering and sample geolocations.

### Estimating effective migration surfaces

Effective migration surfaces were estimated using EEMS [13] and FEEMS [14]. Both models estimate effective migration over a geographic area by comparing genetic similarity and geographic distance among sampled geolocations in the area of interest. In both analyses, estimates are made along a triangular grid, with each sampling location assigned to the nearest vertex on the grid. The grid vertices represent demes in a stepping-stone model, whereby each deme exchanges migrants only with its neighbors (i.e. along the edges connecting the vertices). In cases with equal migration rates between all demes, this produces the null model of isolation-by-distance. Both methods parameterize the effective migration rates (*m*) as deviations from an overall mean rate (µ) across the entire area of interest, so that log_10_(*m*) = 0 is equivalent to the overall mean rate, and log_10_(*m*) = 1 corresponds to effective migration at a rate ten times higher than the overall mean [13].

Effective migration rates in EEMS are estimated using vertex-based Voronoi tessellations, with model parameters fit by MCMC [13]. We used 18 million MCMC iterations with 14 million iterations of burn in for *A. gambiae*, 14 million iterations (10 million burn in) for *A. coluzzii*, and 6 million iterations (2 million burn in) for *A. arabiensis*. The number of iterations was based on observed convergence and accepted parameter distributions (following recommendations in the EEMS documentation) across 10 independent chains per species. For all three species, we thinned the MCMC every 9,999 iterations and set the EEMS grid resolution to the maximum number of demes allowed by the software (1000 demes). For *A. gambiae*, the first 10 independent MCMC chains found separate local optima. We subsequently ran 20 additional independent chains and selected the 10 chains with the highest log-likelihoods (of the 30 chains that were run) to produce the final EEMS output shown in the figures.

Two types of maps are used to display the outputs from EEMS across the area of interest. One shows the posterior means of the effective migration rates, log_10_(*m*). The other map shows exceedance probabilities – in this case, showing where the posterior probability of the log_10_(*m*) being greater than or less than 0 exceeds 0.90 or 0.95. While the former provides a visualization of the point estimates of log_10_(*m*) across space, the latter provides a visualization of where migration rates are most likely to depart from the overall mean rate of log_10_(*m*) = 0.

FEEMS is conceptually similar to EEMS but includes a number of modifications that reduce the computational time by several orders of magnitude [14]. Effective migration rates are estimated for each edge (i.e. the connections between vertices), and model parameters are fit using a quasi-Newton optimization algorithm. A smoothing parameter (lambda) is used to encourage nearby edges to have more similar rates. The main output from FEEMS is a map of the point estimates of effective migration rates, relative to the overall mean rate across the area of interest, similar to the map of posterior means from EEMS. However, FEEMS does not produce an equivalent of the exceedance probability map from EEMS.

To compare the outputs of FEEMS and EEMS, we ran FEEMS with the same 1000-deme grids as above. To assess the effect of grid resolution on model outcomes, with more vertices (demes) than possible in EEMS, we also ran FEEMS using a range of grid resolutions, with edge lengths of 55, 110 and 220 km for each species. To select the smoothing parameter, lambda, we calculated the mean square error from jackknife cross-validation within each set of FEEMS runs. Finally, we visually assessed the resulting maps from each iteration of jackknife cross-validation to assess the robustness of FEEMS estimates to changes at a single deme.

## Supporting information

Supplemental file Ag FEEMS jackknife maps

## Acknowledgements

This study used WGS data produced by the Ag1000G Project (https://www.malariagen.net/project/ag1000g/) with mosquito specimens contributed by 26 partner studies (https://malariagen.github.io/vector-data/studies-ag1000g.html).

## Data Availability

All data analyzed during this study are publicly accessible. Raw sequence reads and sequence read alignments are available on the European Nucleotide Archive under accession number PRJEB42254 (Ag1000G phase 3) and PRJEB2141 (*n =* 165 *A. gambiae* from DRC); and the NCBI Sequence Read Archive (SRA) under accession number PRJNA1159338 (*n =* 5 *A. gambiae* from Malawi). Detailed guidance for accessing sample metadata, raw sequence reads, sequence read alignments, and SNP calls is available at https://malariagen.github.io/vector-data/ag3/ag3.0.html.

