## Supplementary Information for "Variation in spatial population structure in the *Anopheles gambiae* species complex"

McCann *et al.*

### Supplementary Figures

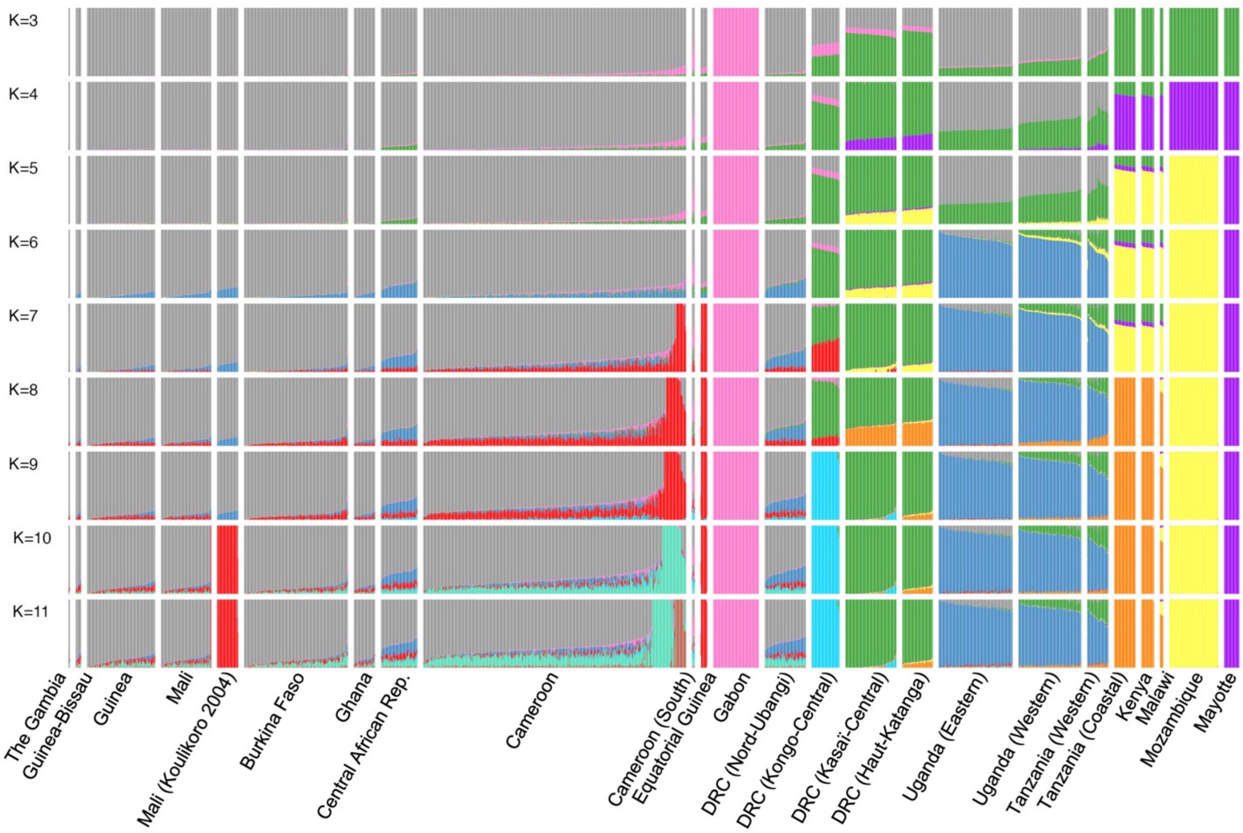


**Fig. S1.** ADMIXTURE with K=3 to K=11 for *A. gambiae*. Each bar represents an individual mosquito, with bars grouped by the country (or region within a country) where the mosquitoes were collected.

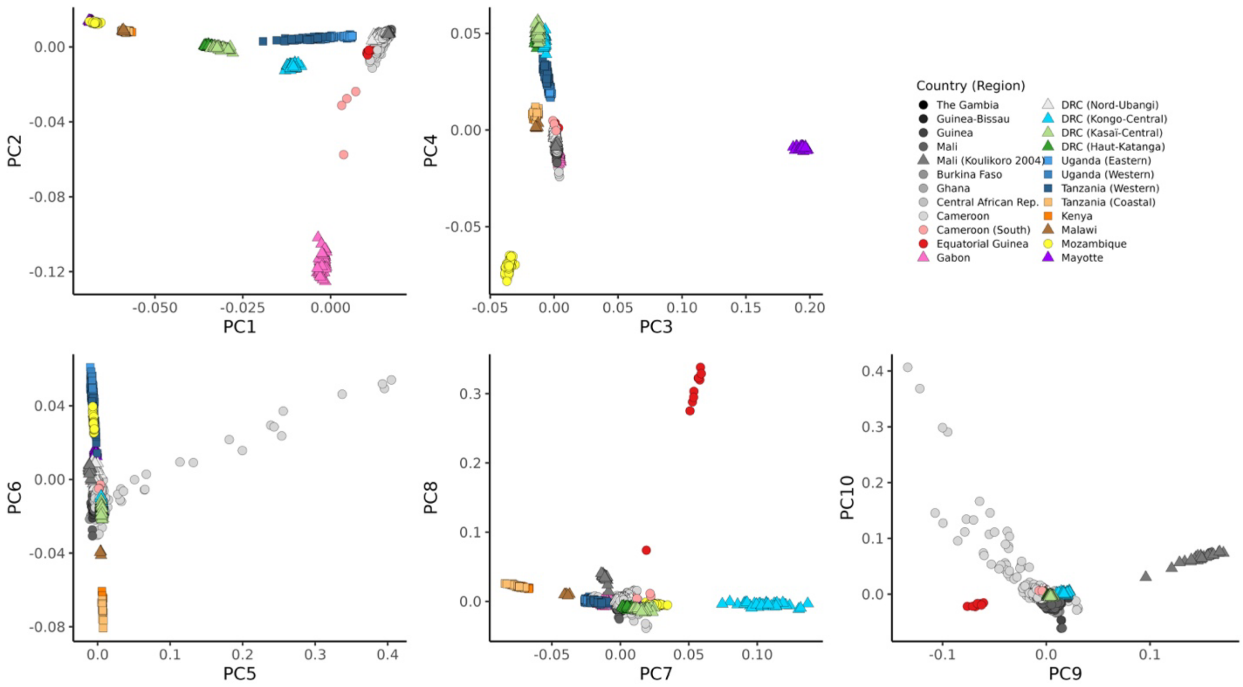


**Fig. S2.** PCA showing PCs 1 to 10 for *A. gambiae*. Each point within a PCA panel represents a single mosquito. Symbol shape and color indicates the country (or region within a country) where the sample was collected.

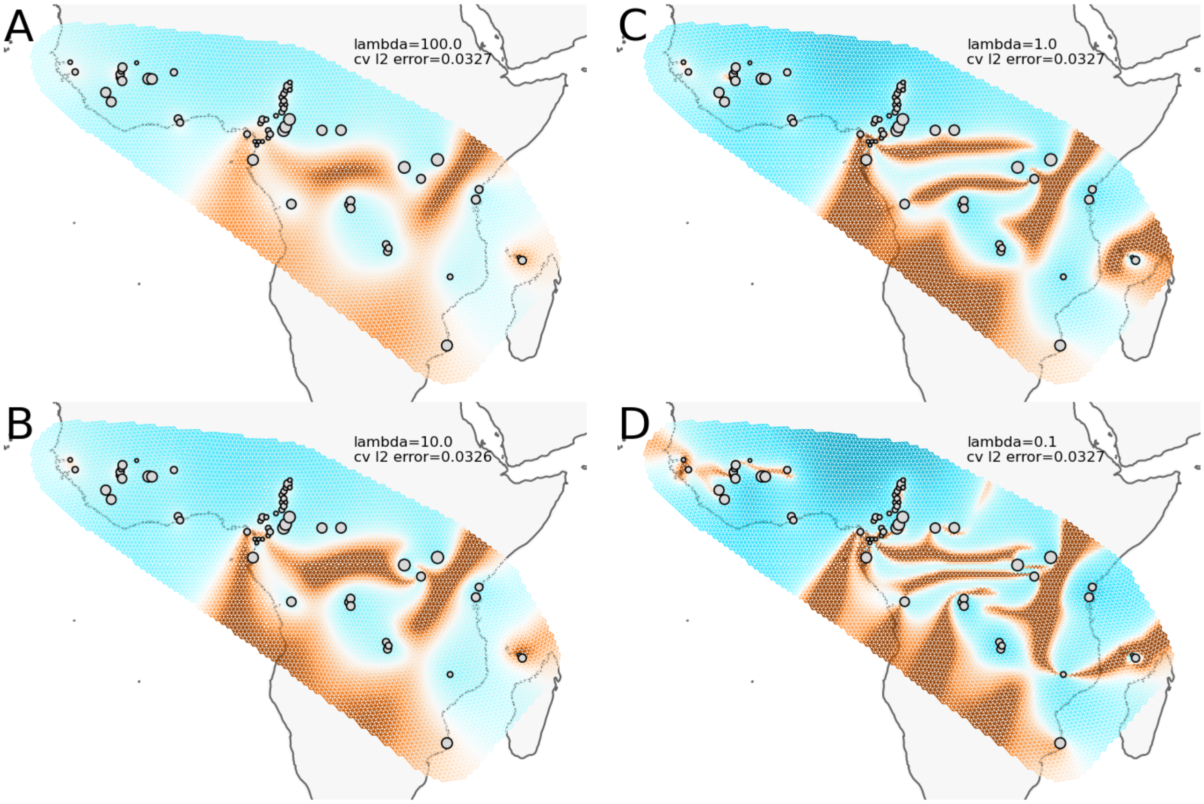


**Fig. S3.** Estimated effective migration surfaces for *A. gambiae* using FEEMS with four different values of lambda and grid edges of about 55 km. All panels use the same color scale for the $\log_{10}$ of the migration rate ($m$). Gray circles show the geolocations of samples included in the analysis, aligned to the nearest node on the grid, with circle size scaled to the number of samples.

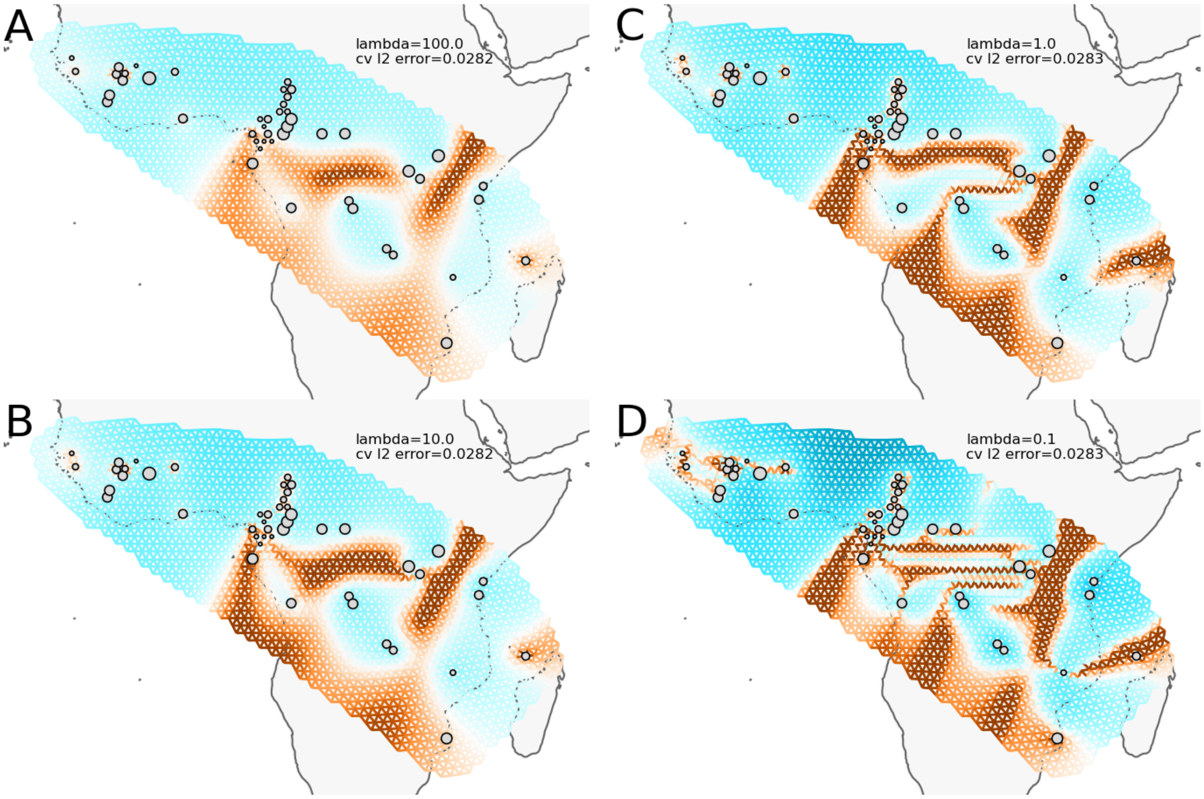


**Fig. S4.** Estimated effective migration surfaces for *A. gambiae* using FEEMS with four different values of lambda and grid edges of about 110 km. All panels use the same color scale for the $\log_{10}$ of the migration rate ($m$). Gray circles show the geolocations of samples included in the analysis, aligned to the nearest node on the grid, with circle size scaled to the number of samples.

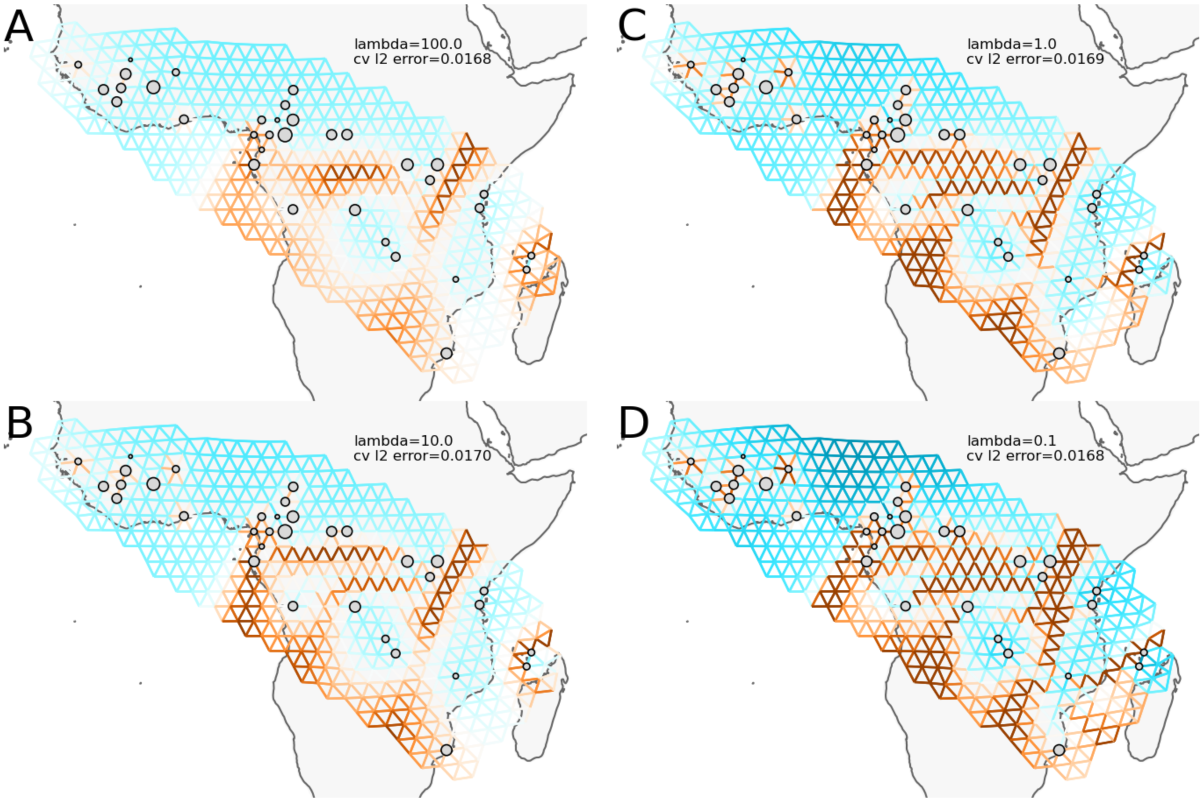


**Fig. S5.** Estimated effective migration surfaces for *A. gambiae* using FEEMS with four different values of lambda and grid edges of about 220 km. All panels use the same color scale for the $\log_{10}$ of the migration rate (*m*). Gray circles show the geolocations of samples included in the analysis, aligned to the nearest node on the grid, with circle size scaled to the number of samples.

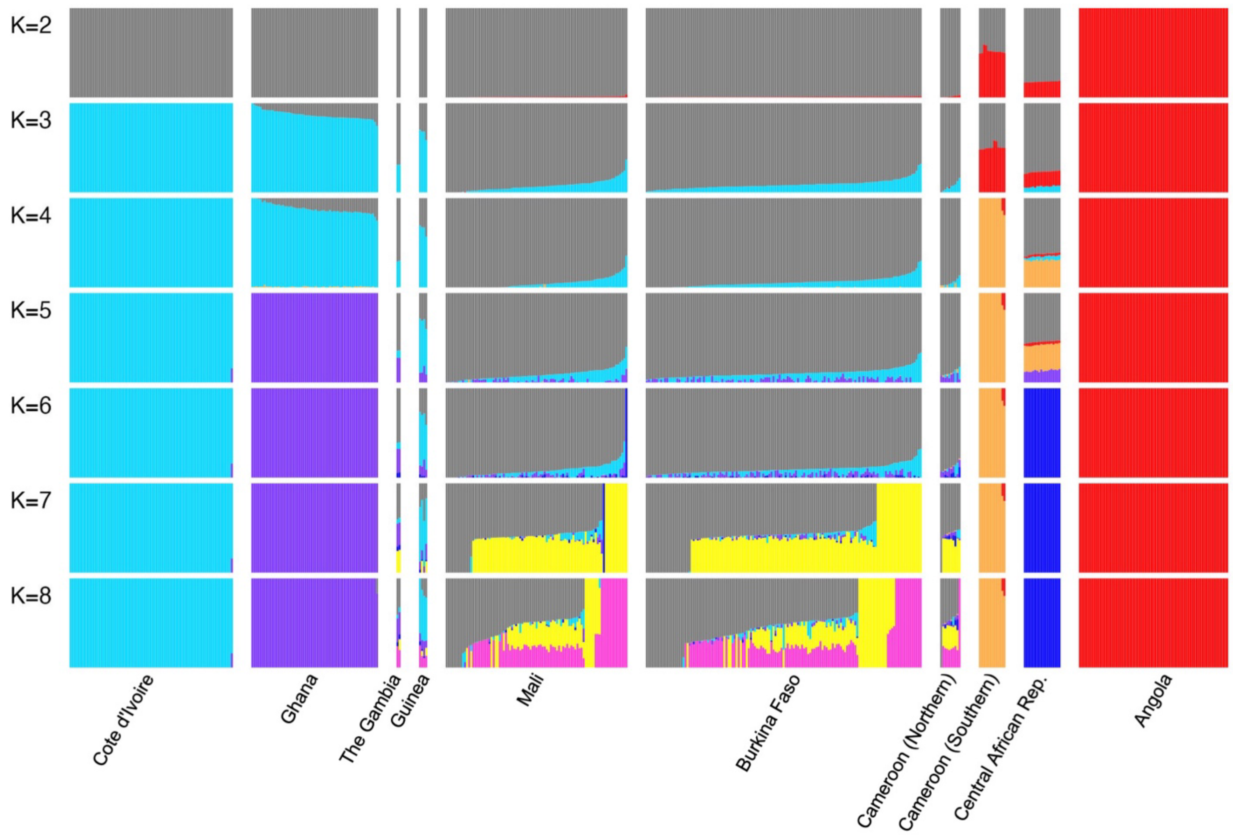


**Fig. S6.** ADMIXTURE with K=2 to K=8 for *A. coluzzii*. Each bar represents an individual mosquito, with bars grouped by the country (or region within a country) where the mosquitoes were collected.

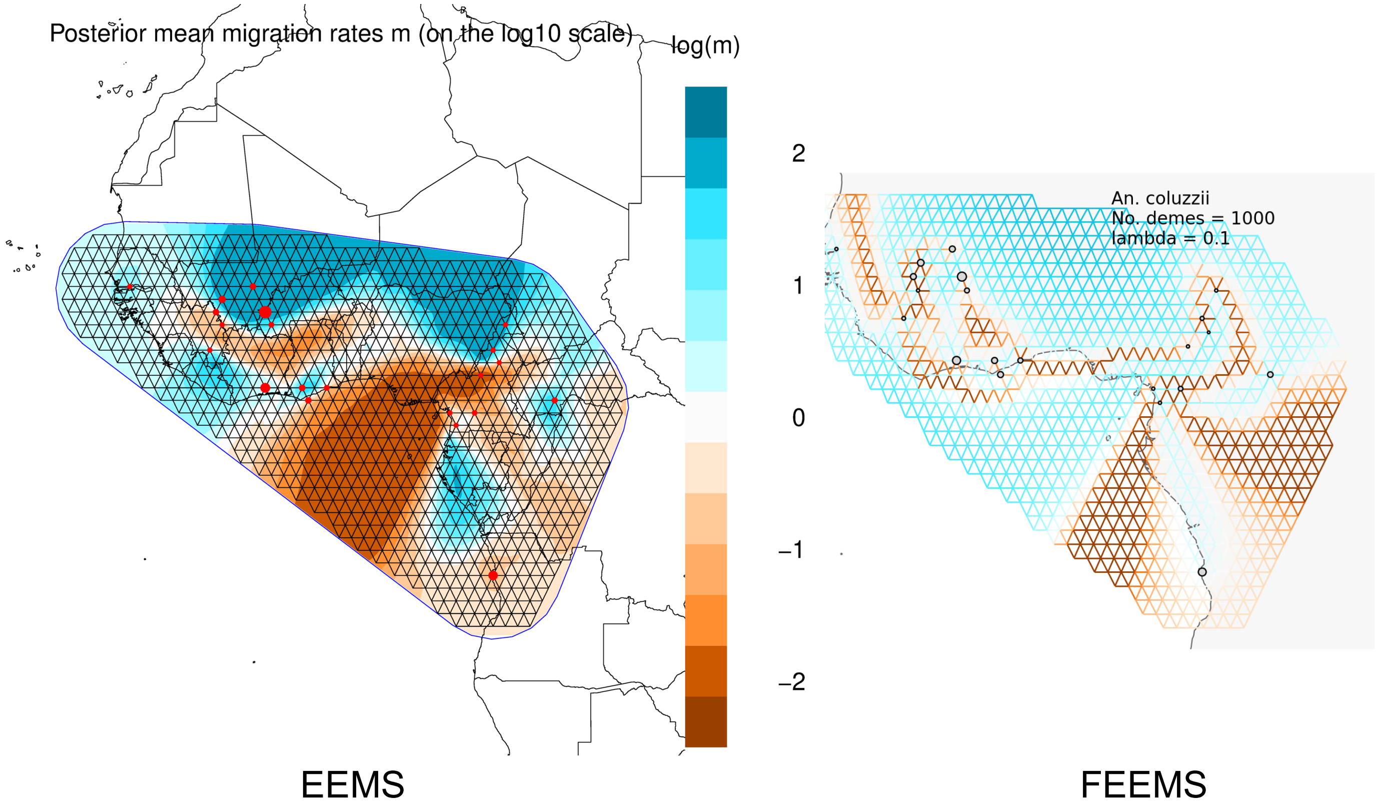


**Fig. S7.** Estimated effective migration surfaces for *A. coluzzii* using EEMS and FEEMS methods. Left panel, output from EEMS. Right panel, output from FEEMS. Both panels show a triangular grid based on 933 nodes (demes) and use the same color scale for the $\log_{10}$ of the migration rate ($m$). Red circles (left) and gray circles (right) show the geolocations of samples included in the analysis, aligned to the nearest node on the grid, with circle size scaled to the number of samples.

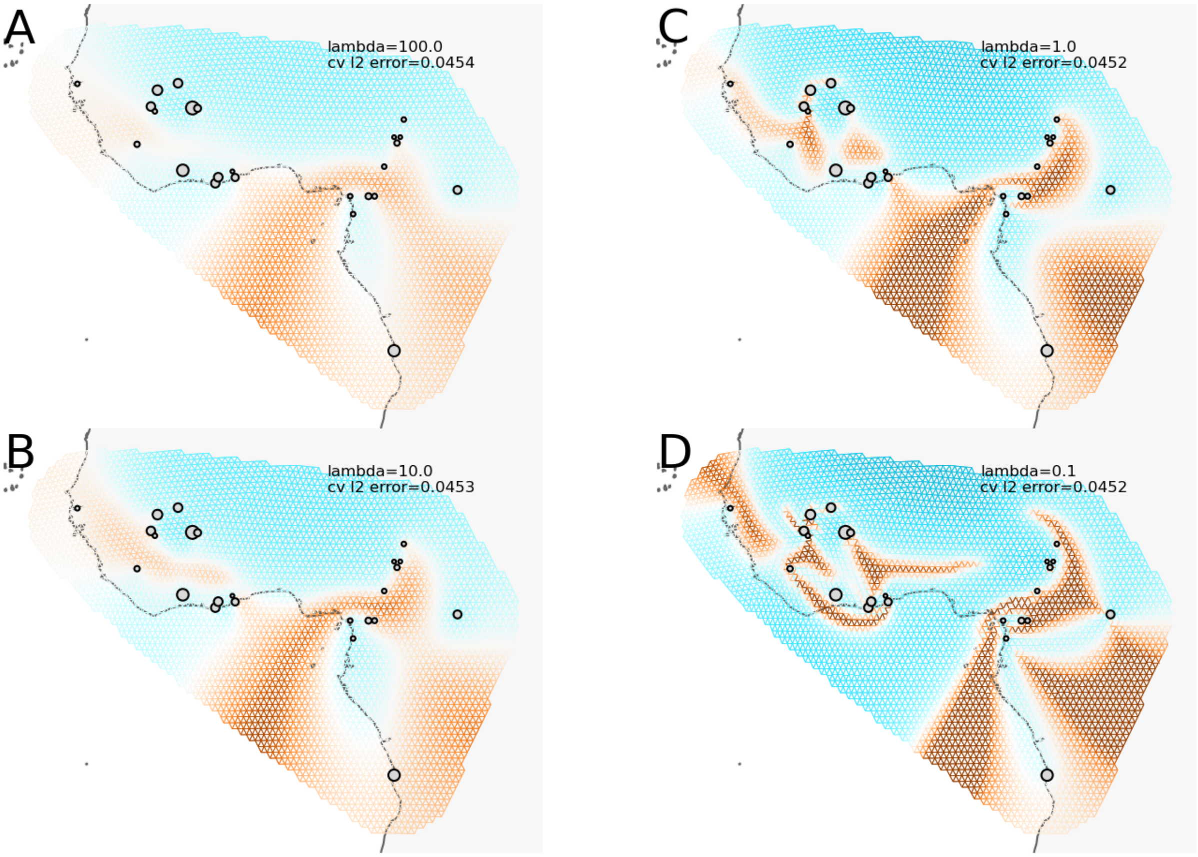


**Fig. S8.** Estimated effective migration surfaces for *A. coluzzii* using FEEMS with four different values of lambda and grid edges of about 55 km. All panels use the same color scale for the $\log_{10}$ of the migration rate ($m$). Gray circles show the geolocations of samples included in the analysis, aligned to the nearest node on the grid, with circle size scaled to the number of samples.

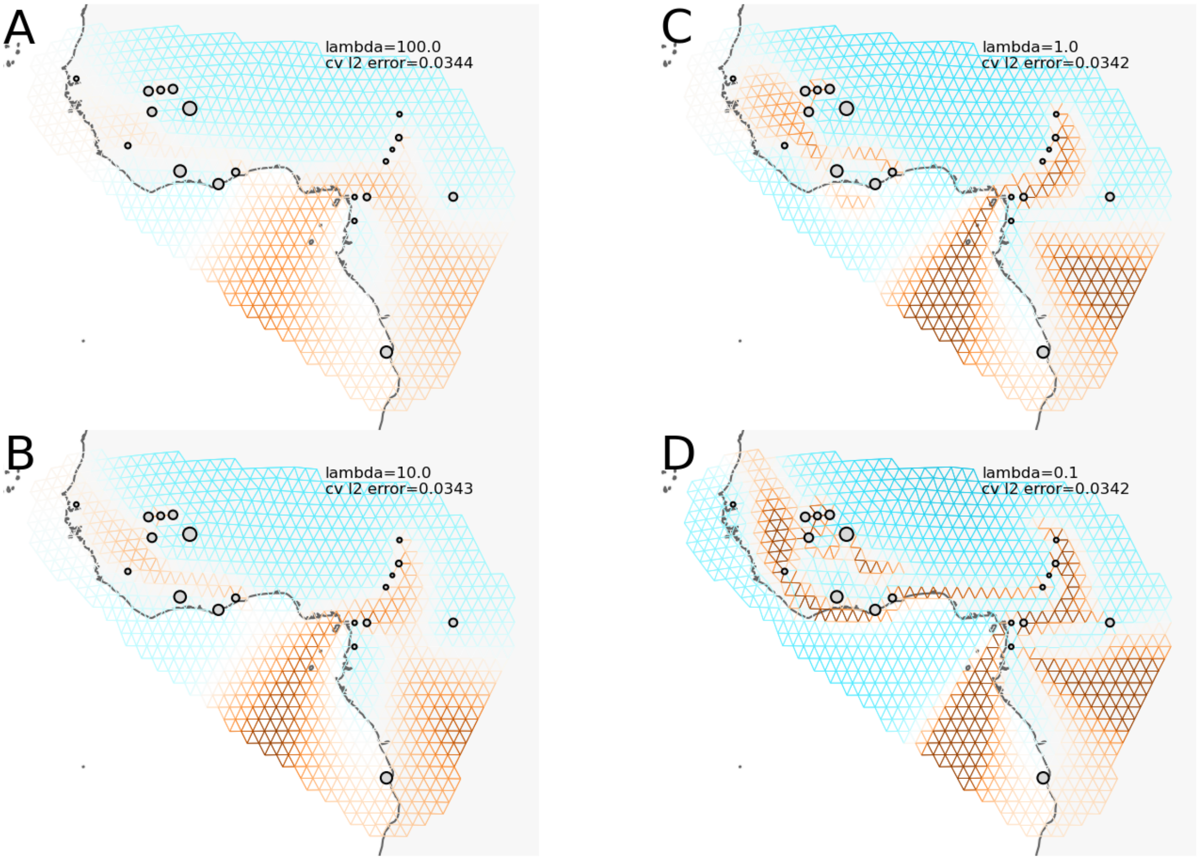


**Fig. S9.** Estimated effective migration surfaces for *A. coluzzii* using FEEMS with four different values of lambda and grid edges of about 110 km. All panels use the same color scale for the $\log_{10}$ of the migration rate ($m$). Gray circles show the geolocations of samples included in the analysis, aligned to the nearest node on the grid, with circle size scaled to the number of samples.

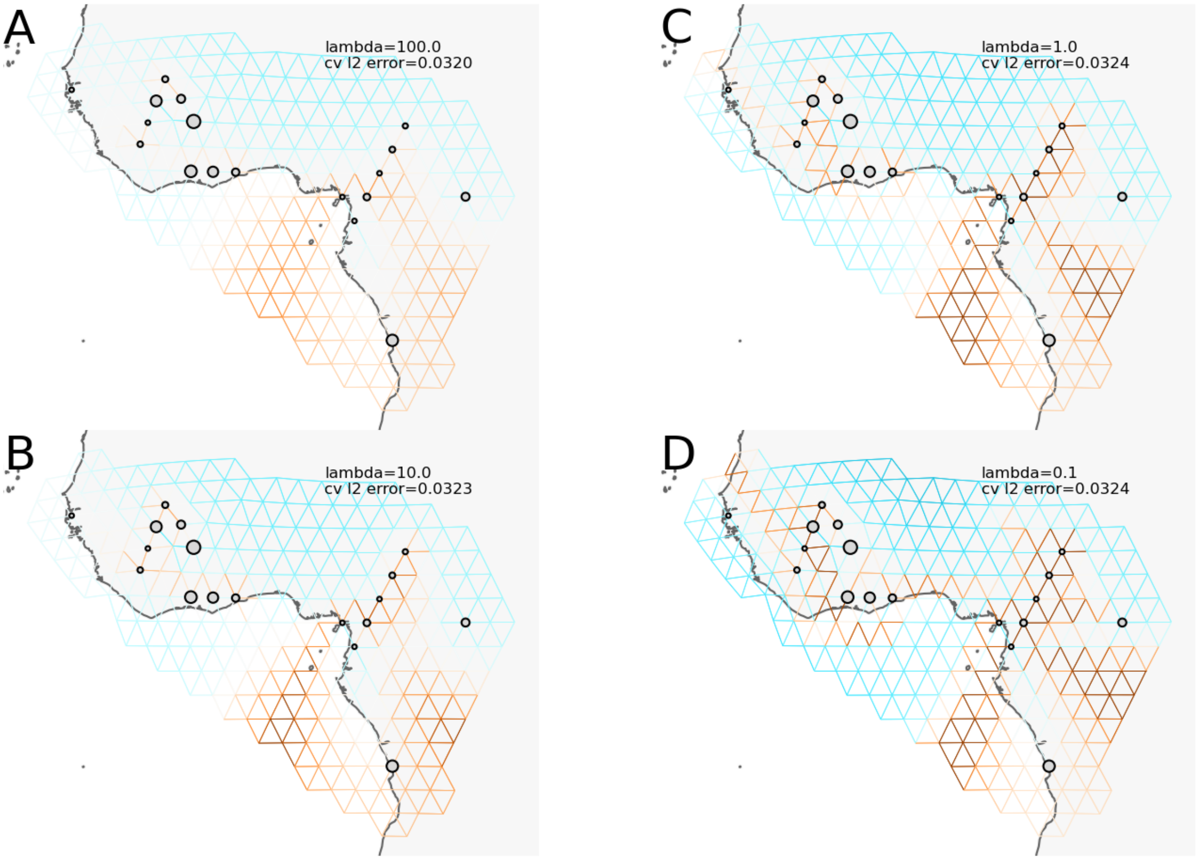


**Fig. S10.** Estimated effective migration surfaces for *A. coluzzii* using FEEMS with four different values of lambda and grid edges of about 220 km. All panels use the same color scale for the $\log_{10}$ of the migration rate (*m*). Gray circles show the geolocations of samples included in the analysis, aligned to the nearest node on the grid, with circle size scaled to the number of samples.

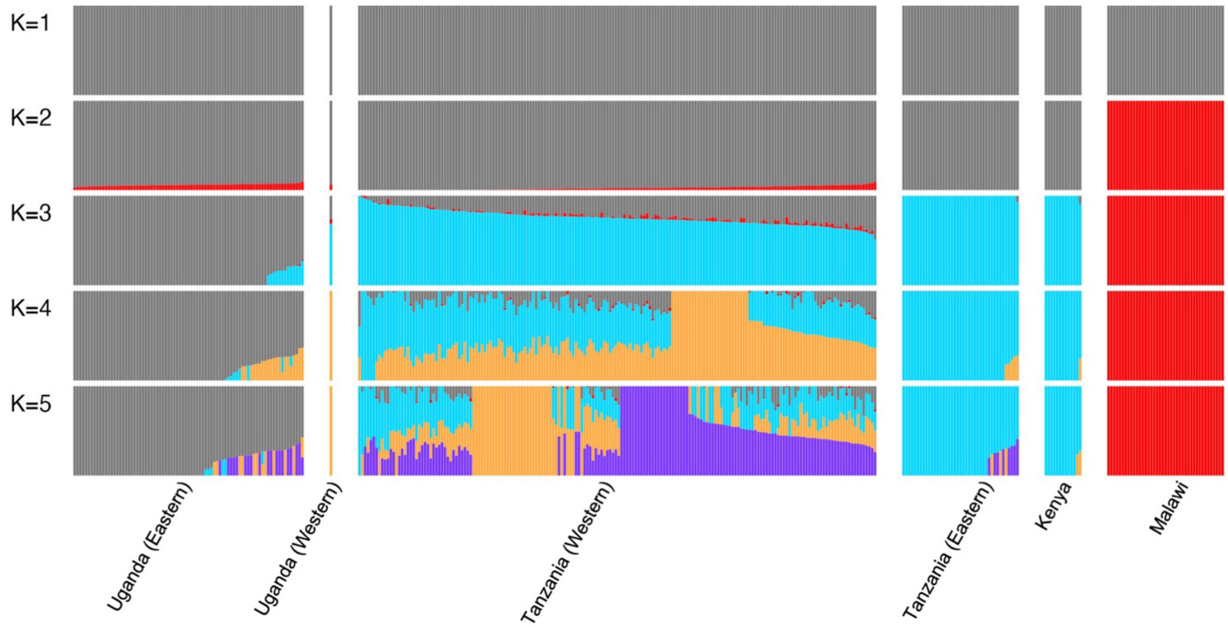


**Fig. S11.** ADMIXTURE with K=1 to K=5 for *A. arabiensis*. Each bar represents an individual mosquito, with bars grouped by the country (or region within a country) where the mosquitoes were collected.

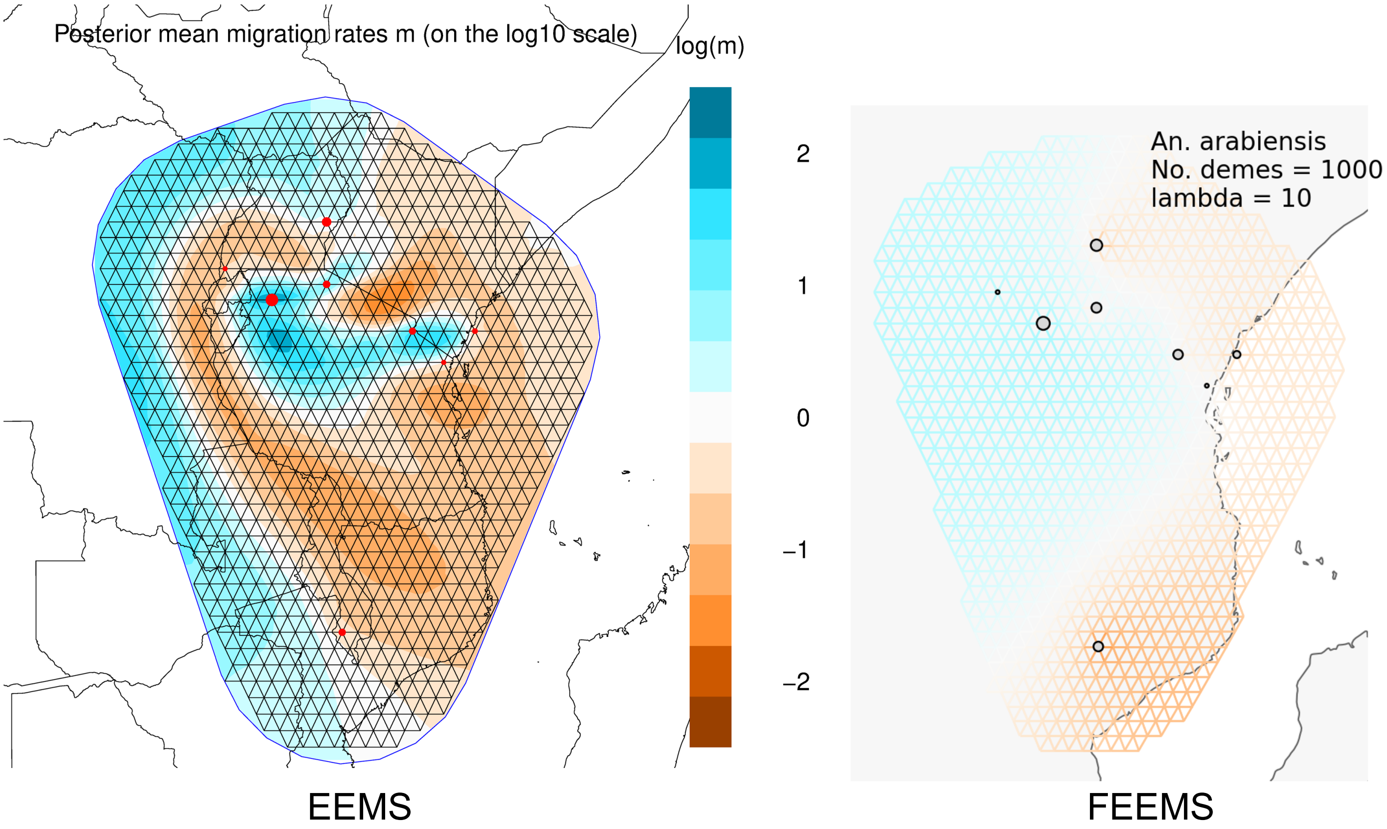


**Fig. S12.** Estimated effective migration surfaces for *A. arabiensis* using EEMS and FEEMS methods. Left panel, output from EEMS. Right panel, output from FEEMS. Both panels show a triangular grid based on 940 nodes (demes) and use the same color scale for the $\log_{10}$ of the migration rate (*m*). Red circles (left) and gray circles (right) show the geolocations of samples included in the analysis, aligned to the nearest node on the grid, with circle size scaled to the number of samples.

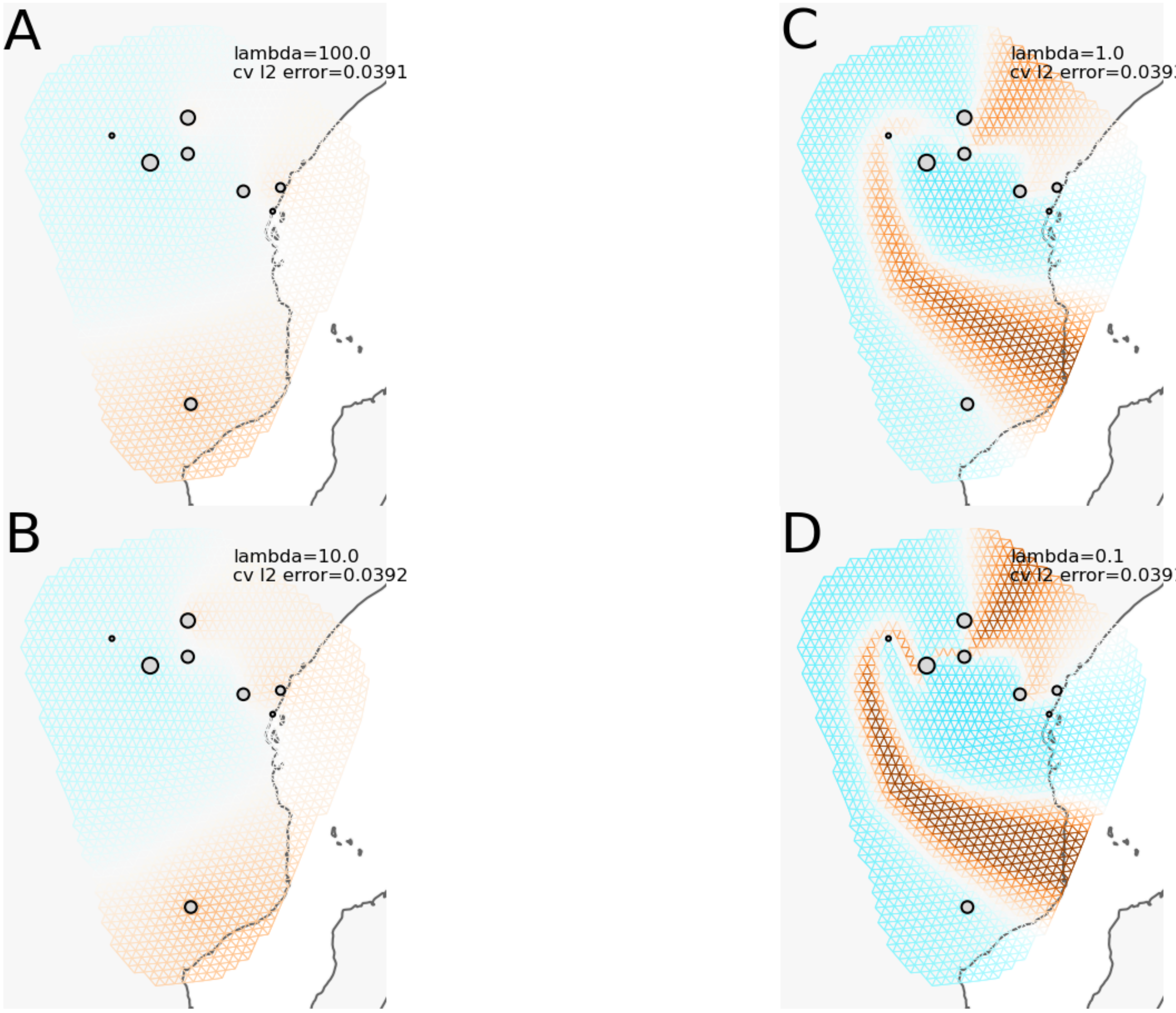


**Fig. S13.** Estimated effective migration surfaces for *A. arabiensis* using FEEMS with four different values of lambda and grid edges of about 55 km. All panels use the same color scale for the $\log_{10}$ of the migration rate ($m$). Gray circles show the geolocations of samples included in the analysis, aligned to the nearest node on the grid, with circle size scaled to the number of samples.

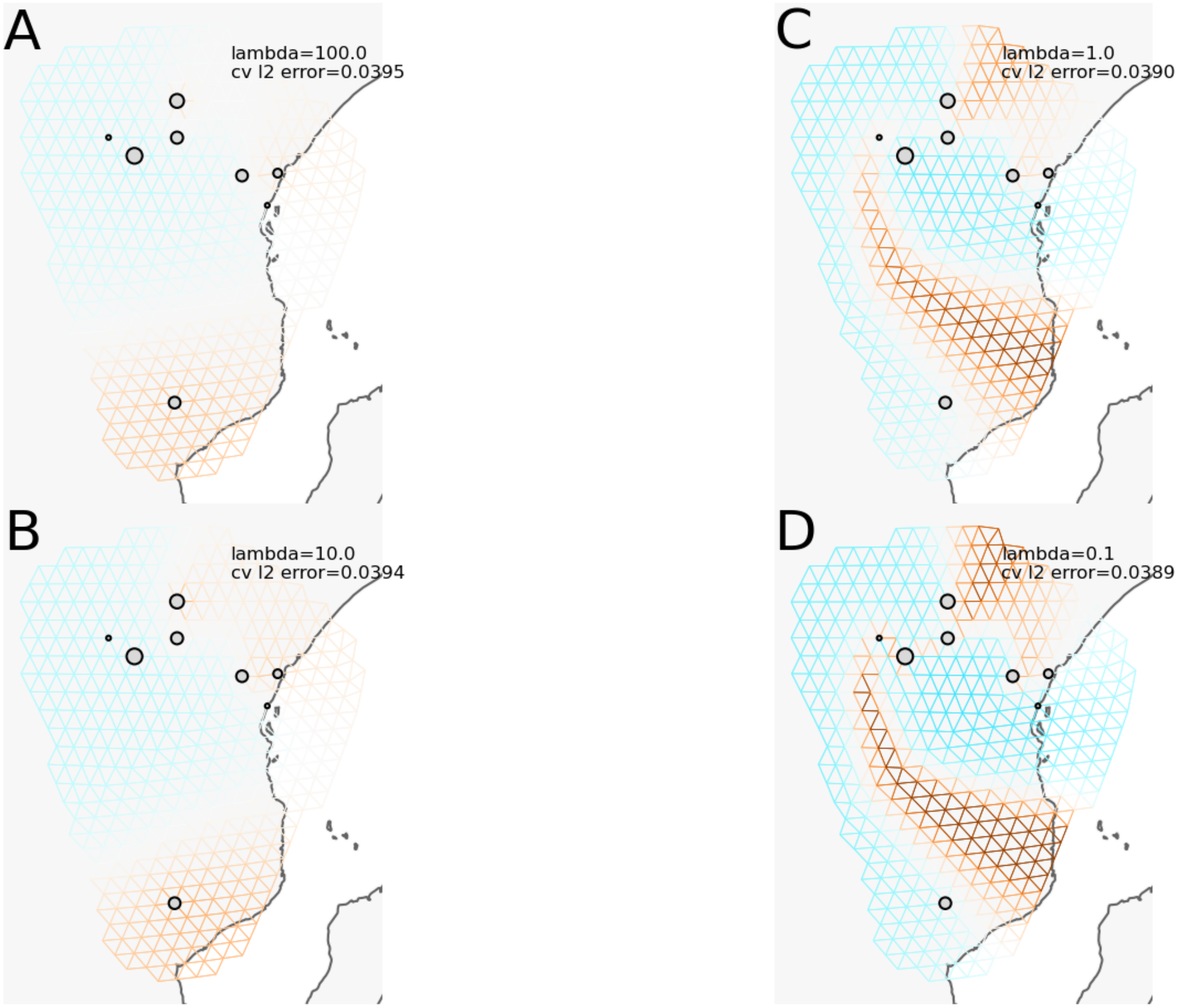


**Fig. S14.** Estimated effective migration surfaces for *A. arabiensis* using FEEMS with four different values of lambda and grid edges of about 110 km. All panels use the same color scale for the $\log_{10}$ of the migration rate ($m$). Gray circles show the geolocations of samples included in the analysis, aligned to the nearest node on the grid, with circle size scaled to the number of samples.

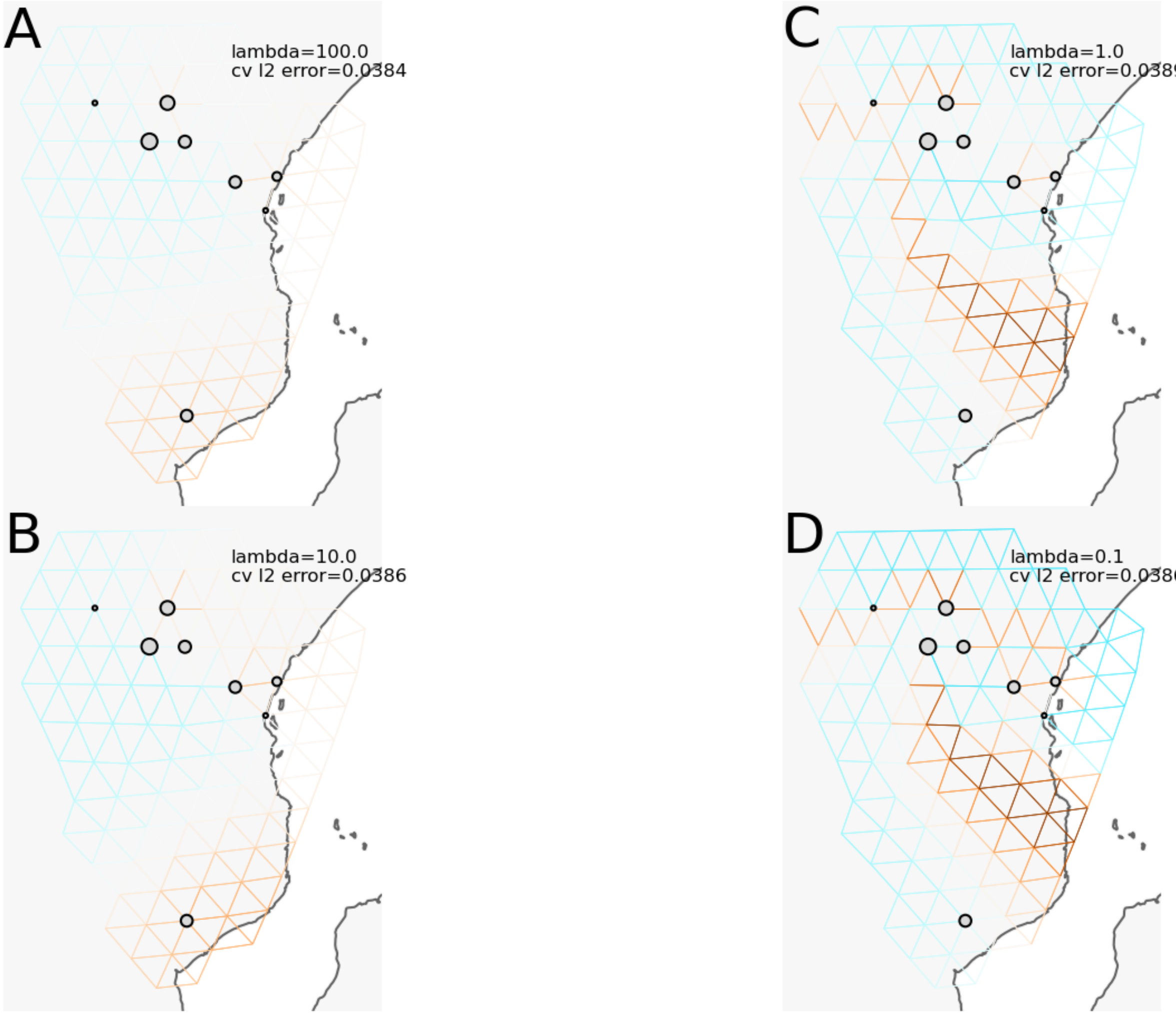


**Fig. S15.** Estimated effective migration surfaces for *A. arabiensis* using FEEMS with four different values of lambda and grid edges of about 220 km. All panels use the same color scale for the $\log_{10}$ of the migration rate (*m*). Gray circles show the geolocations of samples included in the analysis, aligned to the nearest node on the grid, with circle size scaled to the number of samples.
