## Supplementary figures and images for "Variation in spatial population structure in the *Anopheles gambiae* species complex"

### Supplemental file Ag FEEMS jackknife maps

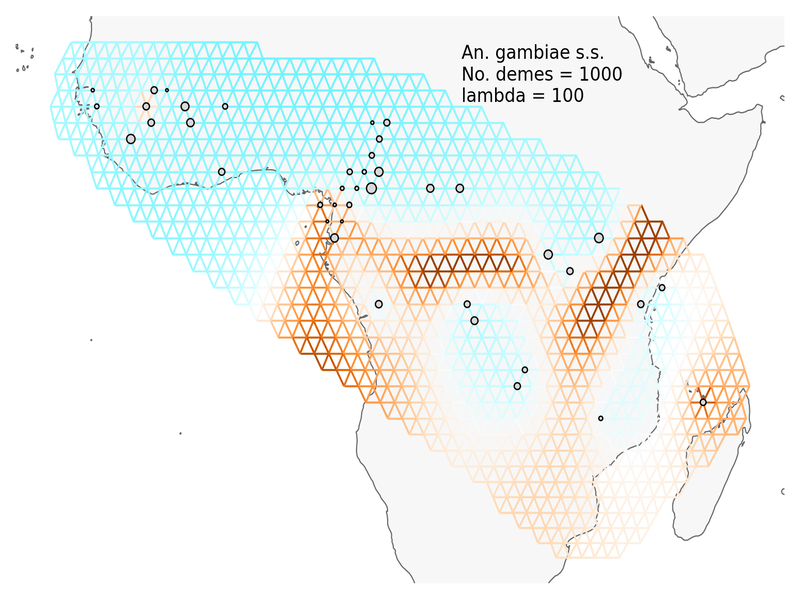
